# Diametric Property of Rotational and Spiral Doppler Effect in Human Heart Atrial Fibrillation

**DOI:** 10.1101/2020.06.21.163642

**Authors:** D Rubenstein

**Affiliations:** Prisma Health

## Abstract

Compass mapping of moving rotors during atrial fibrillation unveiled unique properties of a different form of Doppler phenomenon. New rotational (RDE) and spiral Doppler effect (SDE) equations are derived and explain the diverging frequency changes (increasing and decreasing) that occur simultaneously on either side of passing rotor. These equations confirm that regions of maximal frequency are shifted 90° compared to moving sources that exhibit classical doppler effects. The new SDE equation predicts sequence reversal, side-dependent frequency changes, and an unpaired strong-side wave front strike previously observed in patients with persistent atrial fibrillation. Conclusion: The diametric property of SDE is a mechanism behind dysynchrony of atrial fibrillation when a spinning source of wave fronts moves between two stationary electrodes. The diametric property of SDE in other fields of science are discussed.

**Statement of Significance:** Spinning sources of periodic wave fronts during atrial fibrillation do NOT exhibit classical Doppler effects when they approach and pass between 2 adjacent equidistant electrodes. The wave front is a spinning single spiral arm and frequency of wave front strikes is not only dependent upon distance between successive WF crests, but it is also dependent upon changes in rotation. Rotational and spiral math were developed and to derive new rotational and spiral wave front Doppler effect equations in two dimensions. These equations predict the unique and profound diametric property that have been observed clinically in which the strong side electrodes record an approaching increasing frequency while the weak side electrodes record an approaching decreasing frequency. The solution opens up many new avenues of research, not only into atrial fibrillation, but at all levels of nature, into objects that spin.

## I. INTRODUCTION

A Doppler shift describes a change in the frequency that is sensed by a stationary observer when a source moves while emitting wave fronts at a constant frequency. Christian Doppler in 1842, observed the periodic back and forth shift in light frequency from infrared to visible red that was being emitted from binary stars in rotation (*1*). The higher frequency shift occurred when one star had its maximal angular velocity towards the Earth. This Doppler effect has been confirmed in the longitudinal sound waves and all transverse electromagnetic waves. The Doppler equation is derived on the assertion that periodic waves propagate outward from its source centrifugally.

Therefore, it was puzzling initially to find that a source emitting periodic waves that approached and passed between two electrodes could exhibit both an increasing frequency in one sensor while exhibiting simultaneously a decreasing frequency in the other (*2*). In human hearts with an abnormal heart rhythm, atrial fibrillation, the wave of an electrical impulse through the atrial muscle becomes disorganized in its sequence of activation. There has been mounting evidence that rotors, an action potential (AP) wave front (WF) in the shape of spiral, spins around its central core, and may be a sustaining mechanism of fibrillatory activity (*3-5*). Nearest its central core of a rotor, the single action potential wave front may spin at a fairly constant frequency within a rotation plane that is often parallel to the atrial tissue surface (*6*). It had been previously shown that as a rotor core moved within the atrial tissue, stationary recording electrodes detected a change in the frequency of WFs, and was described (7-9) as a classical Doppler effect (CDE). If the WF propagated centrifugally away from its core, then one might assume that the greatest increase in frequency shift would be detected by electrodes directly in front along the path of the core’s movement. New ablation strategies are currently being formulated to target ablation at atrial regions with the highest dominant frequency of WFs (*5*). Yet, when rotors breached a perimeter of narrow spaced electrodes, the maximal frequency shifts (both highest and lowest simultaneously) occur at either side of the core’s movement (*2*), not directly in front or behind of its path. If local tissue frequencies are used to target sites of ablation (destruction of tissue) as a method to eliminate rotors, then determining which side of that localization becomes important to minimize overall damage.

A cardiac rotor wave front may seem to emit periodic centrifugal propagating waves from its source if observed far away from its core, but a rotor is a single spinning wave that is spiral in shape. When one observes wave front strike frequency from a position close to the center of this rotational source, the wave front crests are propagating not outward, but tangential. The period between wave front strikes is not between successive arms of a spiral but is directly dependent upon the changes in arc length of rotation. The classical Doppler equation cannot be applied. It is because of the patterns of frequency changes now known to occur and that the WF source is a single rotating wave, that different Doppler equations required derivation to be predictive of frequency for this form of WF emission. To solve the spiral Doppler effect (SDE) equation required derivation of a basic rotational Doppler effect (RDE) equation. The new equations accurately predict three main observations of a rotor passing between two electrodes. First, there are diametric frequency changes as observed by 2 equidistant observers placed on either side of a rotors path. Second, there is a sequence reversal WF strikes between the 2 observers. Third, an unpaired wave front strike occurs on the strong side of the path of a rotor. Both new Doppler equations are derived, identifying a special diametric property of frequency changes unique to the observation and recording of spinning bodies in motion. The WF frequency observed is relative to the side of the spin of the moving body. These findings may be at the crux of many other natural spinning or spiral phenomena not yet explained.

### A. Cardiac electrophysiologic waves – Rotor classification

A wave is the disturbance of a field or particles that results in an oscillation which propagates the energy over distances through a mass or medium without the transfer of mass itself (*11*). Waves are typically classified by what media that the waves propagate and in what 3-dimensional orientation that the oscillation occurs. Most commonly observed waves are either longitudinal or transverse, or a combination of the two. A recent addition to the classification of wave types is the rotational wave (*12*). This form of wave rotates around a central position that its wave peak appears to be chasing its own tail. Another visual description of a rotational wave is a transverse wave that propagates in a circle. Physical examples ranging micro to macro scales include circular polarized light to hurricanes and spiral galaxies. Chemical examples of spiral waves first realized with the Belousov-Zhabotinsky reaction (*13, 14*), then within cellular biochemistry in oocyte calcium spiral waves (*15*) and ultimately in electrophysiologic arrhythmias (*16*).

The normal heart beat is a result of an electrical WF, called an action potential (AP) that propagates as a sudden depolarizing change of the cellular membrane voltage, which propagates across the syncytium of cardiac cells. Each cell transduces the electrical event to a mechanical contraction. Thus, the rapid wave propagation creates a single coordinated contraction. Investigative research of this cardiac electrophysiology is vast and the reader is directed to excellent reviews (*17, 18*). Succinctly, pacemaker cells within the sinus node region of the heart exhibit automaticity; the continuous and spontaneous transmembrane voltage change that generates the AP. This is orchestrated by a remarkably intricate set of changes in multiple ion-specific membrane channels that allow ion conductance in a voltage and time-dependent manner. The cycle of sudden depolarization (the upstroke of the AP) and its subsequent slower repolarization repeats about once a second. The AP wave propagates to neighboring cells end-to-end and side-to-side by cellular intercellular gap junction connexin proteins. Electrodes placed on the atrial tissue surface record the exact time that the upstroke of the AP occurs as it propagates directly beneath the electrode. Although conduction of the impulse may propagate anisotropically (*19*), the AP conduction is an aggregate of both longitudinal and transverse propagation.

During a cardiac arrhythmia, the WF propagation path is altered. The propagating AP wave can circle back to where it started, and if it reaches nonrefractory tissue, the AP WF continues to rotate in a circuit-like path or break off into secondary propagating pathways. Here, the upstroke of the AP runs into the end of its own propagation path, aptly named head-meets-tail. The smallest possible rotational path, if stable for several cycles, is termed a rotor (*10*). Rotors have a single AP WF that spiral centrifugally away from an unexcited (non-refractory) central core, called a phase singularity. Rotors may be one of several mechanisms that can sustain the arrhythmia atrial fibrillation for prolonged durations. Experimentally, rotors can be triggered by a single extra AP stimulus in any region of the atria, or by angular interconnection of bundle fibers (*20*). The small core can have an asymmetric shape, have somewhat stable frequency of rotation, and can either meander to other regions of the atrium, or anchor to a single location. The movement of the core, meandering of the rotor, was found to create local WF frequency irregularity and considered a likely cause for disorganized fibrillatory activity (*8*). When a rotor is contributing to WF activity during atrial fibrillation, then longitudinal, transverse and rotational propagation occurs.

In early human trials of localizing rotors during atrial fibrillation, phase mapping methods with a basket catheter requiring marked interpolation between widely spaced electrodes. These potential rotor sites were then targeted for ablation (*21*). More recent mapping and ablation studies utilizing this method have not proven beneficial (*22*). Atrial fibrillation, an arrhythmia that affects millions of people (*23*), causative to marked health care expenditures (*24*), is still poorly understood. In a direct electrical recording method, labelled as compass mapping, we were able to accurately identify the specific rotor sites and time that a rotor meandered past a perimeter of narrowly spaced electrodes (*2*). Rotor activity recorded by this method showed that their existence to be more dynamic, with quite variable durations, and locations. It was a rare occurrence that a rotor was stationary at a single stable location. This method required 3 forms of simultaneous electrode recordings confirming the breach of the perimeter. In the process of recording by this new method, as the rotor passed between 2 electrodes, we discovered a new form of Doppler effect. Even though a single rotor approached 2 narrowly spaced electrodes, one electrode recorded an increasing frequency, while the other electrode recorded a decreasing frequency. This was a highly consistent finding, 54 of 56 transits passed the perimeter. A 2mm distance separated highest frequency of WF strikes to lowest frequency. Such a small distance would confound most direct contact mapping methods without sufficient detailed resolution let alone all non-contact mapping methods.

### B. Side Dependent Cardiac Rotational Frequency Effects

In our detailed study, spiral WFs from rotor activity were tracked as rotors breached the perimeter of the circle (*2*). AP wave frequencies did change, but the frequency changes were shown to be side and spin direction dependent of the rotor. Figure 1 here is a schematic of the results of the actual recordings obtained from multiple simultaneous unipolar and narrow-adjacent bipolar electrodes. A spiral WF (***S***_***Sp***_) approaches the perimeter of electrodes, and the sequence of activation is recorded showing the path of WF propagation (Fig 1, position A). As ***S***_***Sp***_ reaches a mid-position between 2 narrowly spaced electrodes, the WF frequency recordings shows opposite direction of change (position B). At position B, the wave front strike at electrode e_3_ electrode was ½ of a full rotation in time (earlier in phase, thus increased frequency) compared to e_4_ (later in phase, decreased frequency). This method identified where and when along the perimeter that the rotor moved within the perimeter of electrodes. Additional confirmation that the rotor had meandered within the perimeter was made when the unipolar recordings showed sequential activation around the perimeter, thus exhibiting the head-meets-tail wave sequence, as well as by cross-circle recordings of double potentials. One is referred to the Methods section (2). The fact that a single rotor can create simultaneous but diametrically opposite changes in frequency in 2 closely positioned electrodes as the rotor bisects their position, requires the derivation of a new form of Doppler equation to accommodate the side dependency and spin direction with respect to path of motion of the source. This new identified frequency phenomena along with the newly derived equations that support and predict such physical results will from here forward be described as **the diametric property of rotational or spiral Doppler effects.**

**Figure 1.**
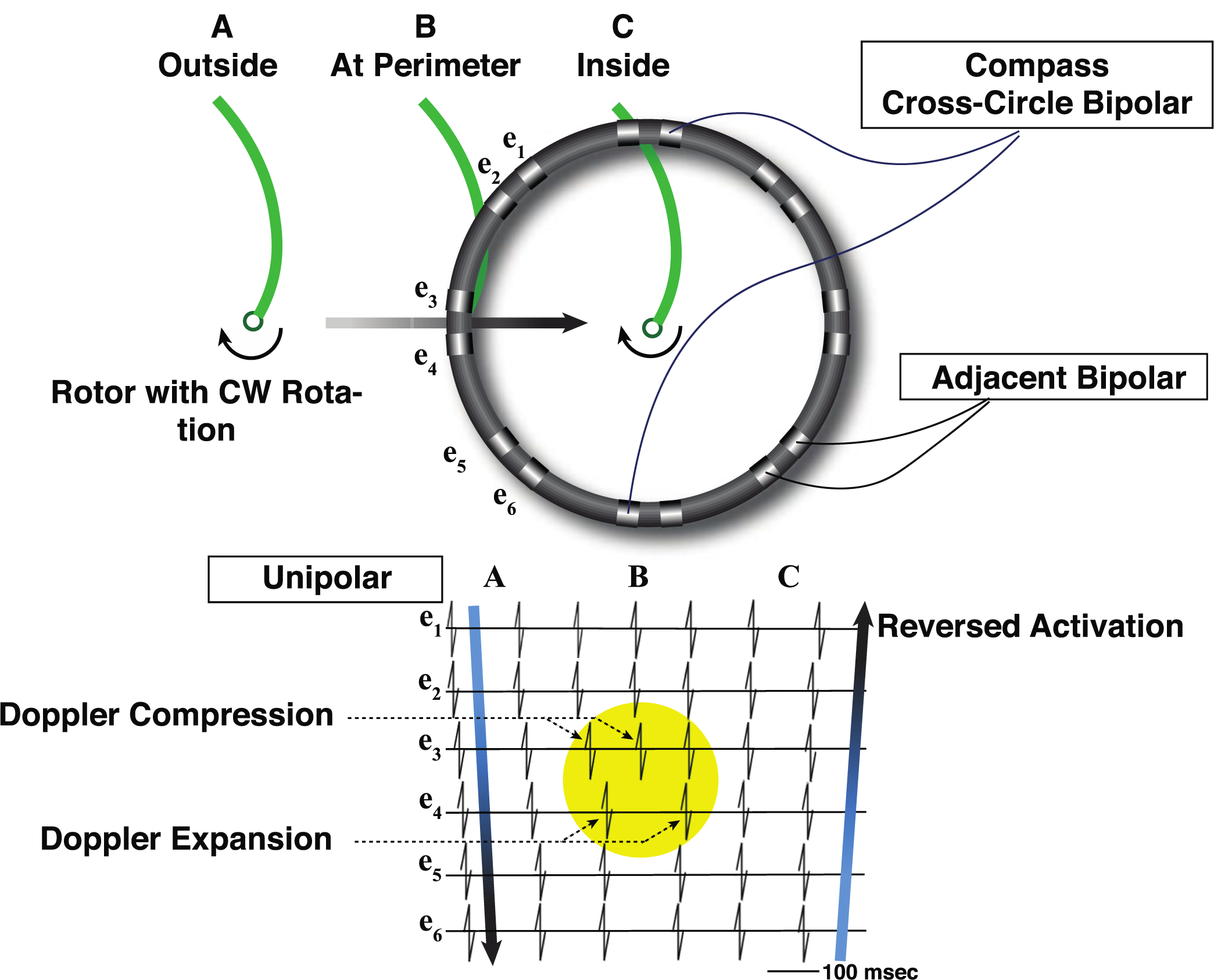
Schematic path of a clockwise (CW) cardiac rotor. Top. The rotor core with an AP wave front (green curved line) moves along a path (gray arrow) between electrodes e_3_ and e_4_ that are a portion of a circular perimeter of electrodes. The rotor has a clockwise rotation. Bottom. Schematic of unipolar recordings of the sequence of the wave front strikes that reach six electrodes (e_1_-e_6_). At position A, the rotor core is outside the perimeter and the unipolar electrodes record wave front strikes sequentially from e_1_ to e_6_. The sweep time for the wave front to pass across all electrodes of the circle perimeter is < 50% of time for one complete revolution. At position B, the rotor core is directly between e_3_ and e_4_ (yellow circle), Unipolar AP WF signals from e_3_ and e_4_ are now separated in time by ½ the rotation (180° out of phase). An increased frequency of wave front strikes at e_3_ (Doppler compression) occurred simultaneous to a decreased frequency (Doppler expansion) directly on the opposite side of the rotor at e_4_. The sweep across all the electrodes consumes 50% of the time to complete one rotation. At position C, the rotor has moved within the perimeter of the electrodes. The activation sequence is now reversed, e_6_ to e_1_. The sweep across all electrode consumes 100% of one rotation (head-meets-tail). The time and location of the perimeter breach was easily identified when evenly split double potentials were recorded by a pair of adjacent bipolar simultaneous with a sudden cross-circle bipolar recording of evenly split double potentials (*2*).

## II. METHODS

### A. The Derivation of a Rotational Doppler Effect

#### 1. Rotor rotation plane and the Rotational Distance Unit

In two dimensions, the AP WF has been shown to be spiral in shape, in three dimensions, the structure of the rotor is described as a scroll wave (*25*). The unexcited core of a scroll wave is extended perpendicularly to the rotational plane to create an unexcited filament. Scroll waves may have different planes of rotation based upon how the filament is aligned with the tissue surface. A transmural scroll wave has its filament extending from the epicardial to endocardial surface. While an intramural scroll wave has its filament parallel to the tissue surface. Tilted plane rotations may also theoretically occur. Supportive evidence for scroll waves has been through computer simulations and animal experimental evidence (*6, 26, 27*). Atrial muscle that line the chamber is thin, only a few mm in thickness. Most meandering transmural scroll waves were found to exist in regions where the atrial muscle was thinnest (*6*). As such, the derivation of equations is in two-dimension but could be expanded to three in later investigation.

A moving source that emits rotational WFs (***S***_***R***_) creates special patterns of frequency changes when observed from stationary positions. The WF of a cardiac rotor, propagates in a circle around its central unexcited core, before extending out in spiral shape. To simplify the understanding of these patterns, an example is presented first with a rotational WF crest having a straight line extending out. A need became apparent to distinguish and separately describe a rotational wave from that of a spiral wave. The methods used to derive Doppler equations of a simple rotational wave first were essential to be able to derive spiral Doppler equations. The new parameters developed here, methods, and graphical presentation might provide the ability to solve frequency related phenomena from rotational or spiral WFs that appear in other forms of nature.

In Figure 2, a ***S***_***R***_ emits a rotational WF of constant angular velocity (***ω***_***s***_) and moves with a constant velocity (***v***_***s***_) directly between 2 observers, or electrodes. A single long clock hand can simulate this rotation extending out radially and can be used to represent a rotational WF with a constant angular velocity. One could further envision thickness to the clock hand such that it not only extends beyond the perimeter of the clock face, but also the edges of the hand extend above and below the plane of the clock face. Figure 2 shows how the timing and sequence changes of a rotational WF strikes that would occur with three different orthogonal planes of rotation. In all three examples, the clock with a constant angular velocity, sits on the tissue surface and is then moved at a constant velocity along the surface plane between two electrodes, e_1_ and e_2,_ are placed on the tissue surface. The clock center moves at a constant velocity, ***v***_***s***_ along a vector path that is a perpendicular bisector of a line that runs between the 2 electrodes. Thus, e_1_ and e_2_ are always equidistant to the center of the source clock. For purposes of orientation and reference, each electrode site has its own clock that is synchronized with the WF source’s clock. Greater radial distance from the center of the source clock must exhibit a proportional increase in the tangential velocity (***v***_***t***_). Therefore, the tangential velocity of the clock hand increases the further away from its center of rotation where

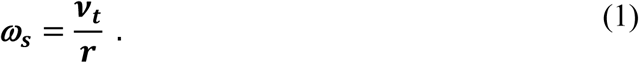

**Figure 2.**
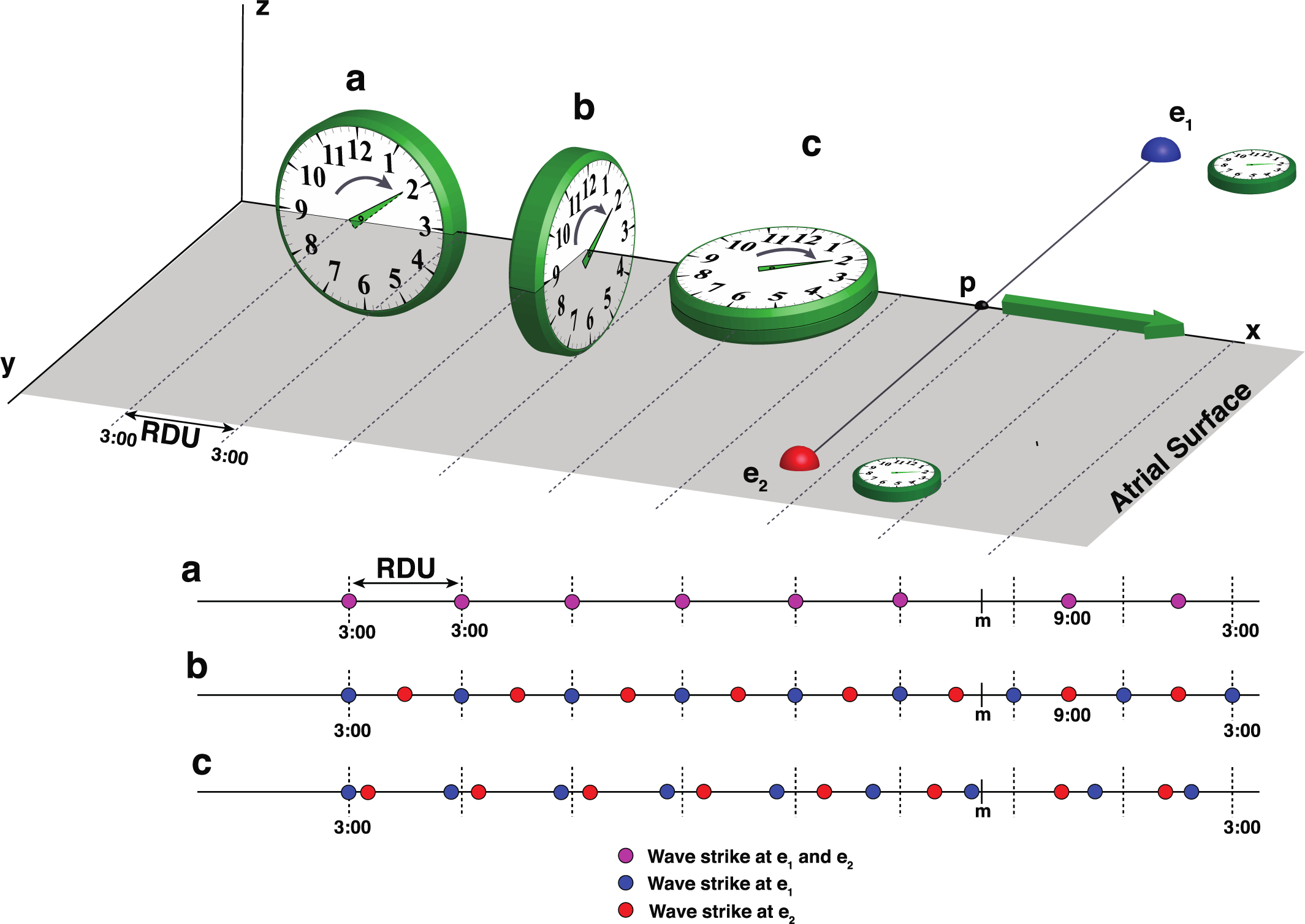
Rotational plane dependent changes in AP wave front strike sequence. Top. Three orthogonal planes of rotation as examples of AP wave front of a rotor spinning in atrial tissue. The green clock hand represents the clockwise rotational AP wave front within the tissue. The gray plane represents the surface of the atrial tissue. Recording electrodes e_1_ and e_2_ are on the tissue surface. The rotor core moves at a constant ***v***_**s**_ along its vector path (green arrow) that bisects a perpendicular line segment at point **p** between e_1_ and e_2_, such that the rotor is always equidistant to e_1_ and e_2_. The distance travelled between each complete revolution is the rotational distance unit (RDU). Bottom. Comparison of rotation plane effects on time sequence of WF strikes at each electrode. Each RDU (vertical dotted lines) is shown as the clock approaches inflection point (**p**). The colored dots correspond to the time at which the wave front strikes the electrode. a) The clockwise rotation plane is in the X-Z plane, with rotation about the Y-axis results in wave strikes at e_1_ and e_2_ to occur simultaneously. When rotor source moves past midpoint, **p**, the wave strike immediately shift 180° in phase. b) The clockwise rotation plane is in the Y-Z plane, with rotation about the X-axis results in wave strikes at e_1_ and e_2_ to occur 180 degrees out of phase. When the rotor moves past **p**, no change in activation sequence is observed. c) The clockwise rotation plane is in the X-Y plane, with rotation about the Z-axis results in sequential activation or e_1_ first followed by e_2_. With each RDU, the time period between WF strikes at e1 shortens, while time between WF strikes at e2 lengthens. When rotor passes **p**, the sequence reverses. Changing the clock rotation to counterclockwise would reverse the sequence changes in the rotation planes as seen in b and c.

In biologic and physical systems, the tangential velocities of WF propagation including action potentials are limited by physical and biophysical restrictions.

A new unit of measurement is presented for calculations and study of frequency changes resulting from any rotational or spiral WF. A rotational distance unit (an RDU) is defined as the distance travelled at a constant source velocity (***v***_***s***_) as the WF makes one complete revolution around its center,

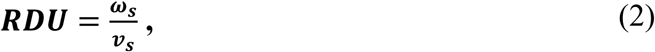

where the fraction is reduced to having units of 2π/distance. This initial essential step with reduction to 2*π* in the numerator, and since ***ω***_s_ = 2*πf*,

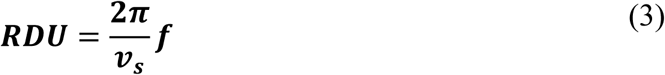

or

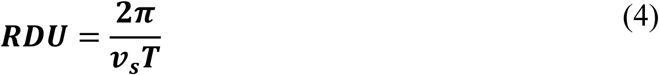

and replacing ***v***_**s**_**T** with D, the distance travelled, then the denominator simplifies to

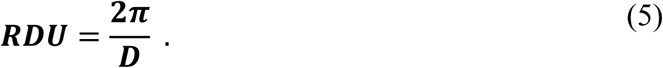

The RDU allows construct of other units, parameters, and graphical presentations that will allow the accurate computation of WF strikes, and frequency calculations at any position of electrodes (or observers). The standard definition of period (***T***) is used as the time between WF strikes at a specific electrode.

In the first example (Figure 2a), the rotational plane is perpendicular to the atrial surface with the rotational axis perpendicular to the line of motion between the two electrodes (analogous to a rolling tire). As the clock moves down its path, with each RDU, the WF strikes electrodes e_1_ and e_2_ simultaneously at precisely 3:00 (pink dots). Once the core has passed by the midpoint (**p**), the AP WF strikes the electrodes at 9:00 with each RDU. In the second example (Fig 2b), the rotational plane is again perpendicular to the atrial surface, but the rotational axis is parallel along the vector path of motion (analogous to a rotating screw, or circular polarized light). Here, at each RDU, the rotational WF strikes electrodes e_1_ and e_2,_ sequentially (blue and red dots), 3:00 and 9:00 respectively, before and after the center of the clock passes the **p**. The sequence of WF strikes between e_1_ and e is 180^0^ out of phase. In the third example (Fig 2c), the rotational plane is parallel to the atrial tissue surface with the rotational axis that is perpendicular to that plane similar to most cardiac rotors as the move across the tissue surface. When the source moves along it path, closer to e_1_ and e_2_, the source WF will strike e_1_ earlier in time by a shortening of the arc length of the source’s clock.

The change in arc length is a change in the angle of the clock governed by the equation

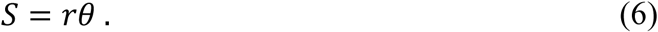

As arclength shortens with subsequent WF strikes at e_1_, the clock hand must sweep further around than the original arc length for the WF to strike to occur at e_2_. With each subsequent complete revolution (RDU) of the source, up until and after the source reaches the inflection point (point **p**), the arc length on the source’s clock shortens for WF strikes at e_1_ and lengthens for WF strikes at e_2_. As the clock approached, the change in the arc length results directly to a proportional change in the angle of the clock hand, resulting in a decreasing period (increased frequency) of WF strikes at e_1_ and an increasing period (decreased frequency) of WF strikes at e_2_. As the core passes the midpoint **p**, the sequence of WF strikes reverses, e_2_ precedes the WF strike at e_1_. Importantly, in this last example, had an observing electrode been placed directly in front of the moving source (at inflection point **p**), WF strikes at that position would have occurred at 3:00 with each RDU as it approached, then at 9:00 after passing **p**. Thus, no change in period, no change in frequency with approaching or receding from point **p**, just an instantaneous phase shift in ***θ*** by 180 degrees or by *π* radians.

#### 2. Clock Hands, Strong and Weak Sides, and Rotational Math, and the derivation of the rotational Doppler equation

##### a. Clock hand as a rotational wave

In Figure 2 example c above, represent the frequency changes observed in the moving cardiac rotor. The profound diametric result that an approaching source emitting periodic rotational WFs exhibited simultaneous but opposite frequency effects, is a side-dependent phenomenon. The diametric property requires one to define the side dependency such that an electrode or an observer sits on a strong side or the weak side to the source. The strong side identifies the side that receives a greater frequency of WF strikes.

To quantitatively understand the diametric property observed, a hypothetical timing experiment is presented (Fig. 3 and 4) where a ***S***_***R***_ passes between 2 equidistant electrodes or observers. The ***S***_***R***_ is represented as a highly accurate clock. The clock is placed flat on the ground with 12:00 aligned to the North. The sweeping clock hand represents the WF crest that extends far out beyond the perimeter of the clock and moves as a single wave. The clock moves along a vector path (***v***_***s***_) to the East (the 3:00 line). Two observing electrodes (e_1_ and e_2_) each have clocks that are precisely synchronized to ***S***_***R***_, are also each placed flat with 12:00 positioned North. The center of each clock represents the exact location of S, e_1_, and e_2_. The clock hand angle, ***ψ***_***c***_, (Figure 3(a)), is measured from 0 degrees placed at 3:00 (such that 2:00 and 4:00 represent +*π*/6 and -*π*/6, respectively). The angles increase positively and negatively until they meet at 9:00 (+*π* and -*π*, respectively). A CCW rotation would reverse the designation of + and – angles from the 0-degree position. Electrodes, e_1_ and e_2,_ are specifically positioned, on both sides and equidistant to the source’s vector path of motion. The vector path of ***S***_***R***_ is a perpendicular bisector between e_1_ and e_2_ along its entire vector path, allowing direct comparison of frequency shifts on either side of a moving and rotating source.

**Figure 3.**
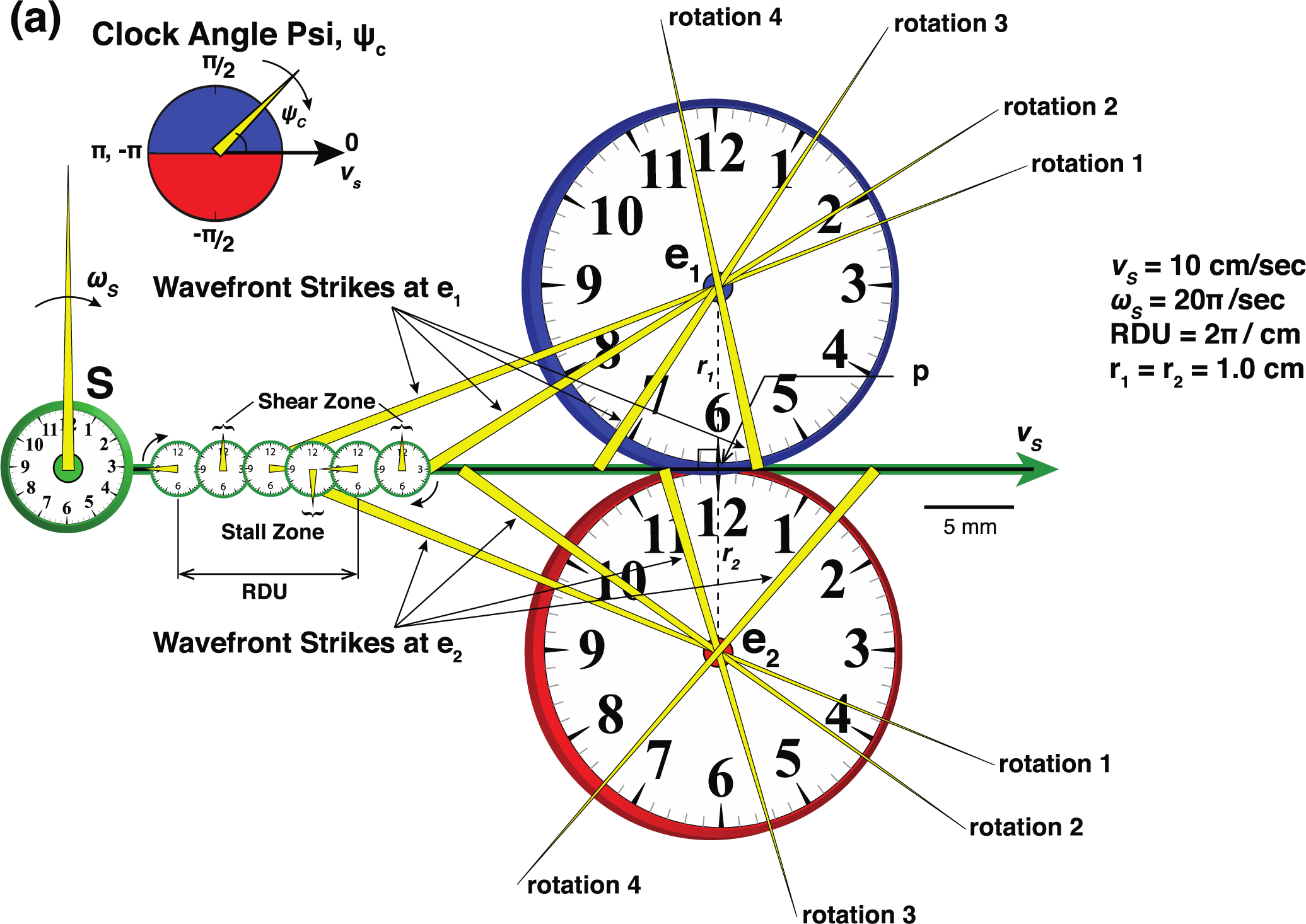
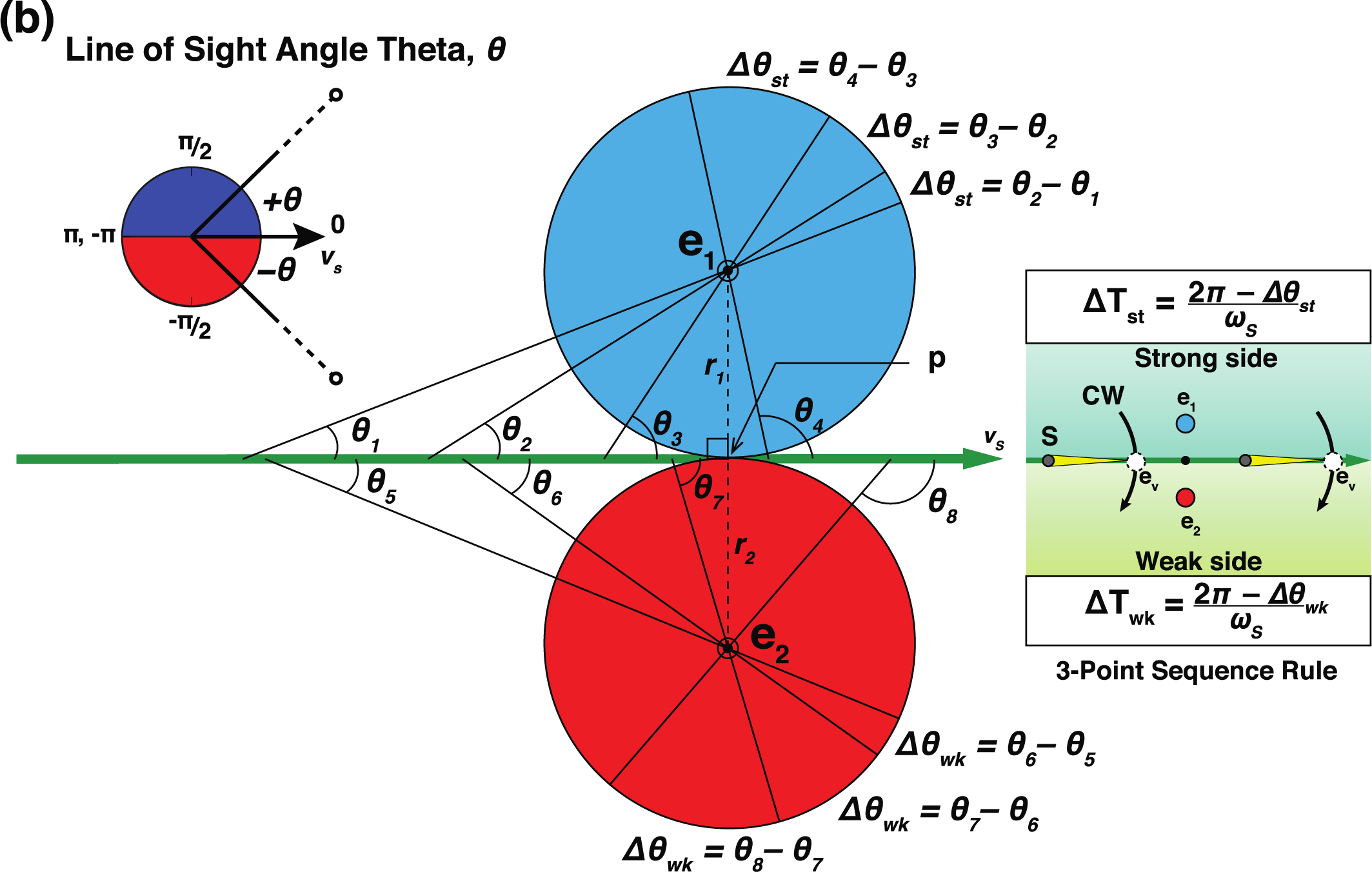
Orientation of Clock Angle *ψ*_*c*_ and Line of Site Angle *θ*. (a) Clock Angle ***ψ***_***c***_. Upper inset shows angle 0° is set at forward vector path at the 3:00 position. During CW spin, the clock hand (thin yellow triangle) of the source (S), is initially positioned at 9:00 (*π*). The source’s clock hand rotates with constant ***ω***_***s***_ at 10 revolutions/sec (20*π*/s). The clock hand is a positive angle at *π* and decreases as ***ψ***_***c***_ rotates towards 3:00 passing +*π*/2 set at 12:00. When clock hand passes 12:00, fastest forward tangential velocity is parallel to ***v***_***s***_ that may create a potential WF shear zone at greater radii. Once the clock hand passes 3:00 (0 radians), ***ψ***_***c***_ is a negative angle as it rotates back towards 9:00 (-*π*), passing -*π*/2 set at 6:00. As the clock hand passes 6:00, the slowest forward velocity is parallel to ***v***_***s***_ that may create a potential WF stall zone at greater radii. The RDU (small clocks along vector ***v***_***s***_) is shown as the clock hand makes one complete revolution. Shear and stall regions would be expected to occur with each revolution at same respective ***ψ***_***c***_, separated by distance D of the RDU. Electrodes e_1_ and e_2_, depicted at the center of synchronized clocks, are positioned 1 cm on either side of the vector path of the source that moves at 10 cm/s. The vector path is a perpendicular bisector between e_1_ and e_2_. Four subsequent WF strikes are shown at each electrode as S moves passed the inflection point **p**. (b). Line of Sight Angle ***θ***. Upper inset shows that angle 0° is set directly along the forward vector path of the source. The positive and negative angle ***θ***s are set to align with the positive and negative angle ***ψ***_***c***_ as seen in 3A. Note that once ***S*** passes the inflection point ***p***, LOS angle ***θ*** is obtuse. The change in angle ***θ*** (**Δ*θ***) between subsequent WF strikes is a positive value at a strong-side electrode, and negative value at a weak side electrode.

**Figure 4.**
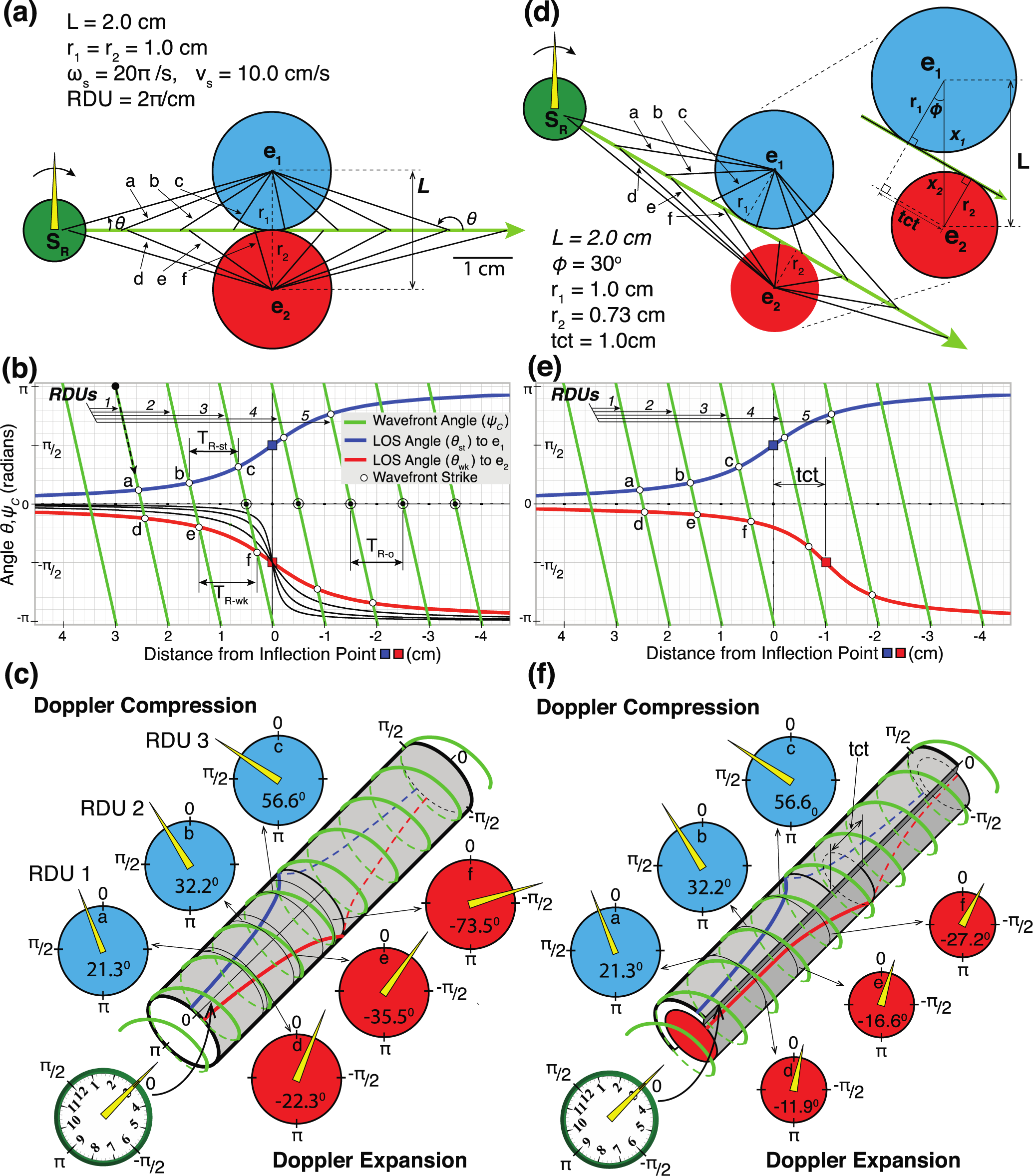
Electrodes on either side of *S*_*R*_’s vector path reveals the diametric property of RDE. (a). Source’s clock, and electrode clocks are synchronized, aligned, and lay within the same horizontal plane. Clockwise ***S***_***R***_ with constant angular velocity 20*π*/sec, moves at constant velocity 10 cm/sec (RPD = 2*π*/1cm) along a vector path which is a perpendicular bisector of line e_1_e_2_ (2.0 cm apart). The electrodes e_1_ and e_2_ are equidistant to ***S***_***R***_ along its entire path of motion. Clocks at e_1_ and e_2_ constructed with each radius 1.0 cm touch each other at the common tangent point **p.** The source clock reads 9:00 when the source is 3mm away from point **p.** One complete revolution of the clock occurs every 1 cm (RDU = 2*π*/1cm) as the source moves toward the electrodes. The clock hand (yellow triangle) extends beyond the perimeter of the clock, rotates and strikes e_1_ before striking e_2_. The first 3 RDUs result in WF strikes at e_1_ at line a,b, and c, and at e_2_ at lines d,e, and f. The distance from **p** and the angle of the WF strikes are listed in table 1. The clock hand strikes the e_1_ clock earlier in its rotation (strong side) while striking e_2_ later in its rotation (weak side). See Supplementary Materials Figure 4 Data File. (b). Graphical presentation of both f(x) = ***ψ*** _***c***_ of the clock hand angle and f(x) = ***θ*** for the LOS angle to both electrodes as ***S***_***R***_ approaches and recedes from the point **p**, the closest approach to electrodes. Note that x-axis is reversed, having increasing positive values going leftward. The LOS angle ***θ*** from the source to e_1_ and e_2_ is calculated by the arccotangent for the strong side (blue curve) and the arccotan curve for the weak side (red curve) from Eq.9 and Eq. Utu10 respectively. Starting point for first RDU (black dot with dashed arrow), begins with the source clock hand at 9:00 (*π* radians) with the source 3 cm away from point p. The RDU lines of the clock angle ***ψ*** _***c***_ (green lines) intersect the arccot curves for the LOS ***θ*** at the moment of WF strike at e_1_ and e_2_ (white circles). Intersection points at a-f correspond to the distance from **p** and angle shown in 4A above and listed in Table 1. RDUs #1-3 show WF strikes at a-c at e_1_ and d-f at e_2_. Arccot curves are shown for comparison if e_2_ sits closer to the inflection point at distances ½, ¼, and ⅛ of r_2_ (black curves). (c). 3-D cylindrical polar plot of fig 4(b), wrapping the graph into a cylinder with radius 1 cm. The clock hand angle rotates around the cylinder (green coil). Distance between coils is the RDU. Electrode e_1_ sits on the strong side of the source and subsequen*t* slices (blue circles) show progressively earlier times of strikes, exhibiting a Doppler compression. This occurs while e_2_ sits on the weak side of the source and subsequent slices show progressively later times of strikes, exhibiting a Doppler expansion. Slices of the cylinder are pulled out to show when the clock hand WF strikes at electrodes e_1_ (blue circles a,b,c) and at e_2_ (red circles d,e,f). (d). Similar to A, the ***S***_***R***_ clock, and electrode clocks are all synchronized, aligned, and lay flat within the same horizontal plane. The vector path of the source is at a tilted (*ϕ* = 30°) compared to A. The path is created such that e_1_ is still 1 cm from the vector’s path of closest approach. The inset of the enlarged electrodes shows the trigonometry used to calculate the transverse common tangent line segment length (tct) and the e_2_ distance to the closest approach of the source (r_2_). Electrodes e_1_ and e_2_ remain at their same position with L =2.0 cm of separation. The tct is 1.0 cm (tct = L sin *ϕ*). The distance (r_2_) of e_2_ to the point of closest approach of the ***S***_***R***_’s vector path is calculated at 0.73 cm (r_2_ = x_2_ cos *ϕ*). (e). Similar to B simultaneous plots of f(x) = ***ψ*** _***c***_ of the clock hand angle and f(x) = ***θ*** for LOS angle to both electrodes. The plot of ***ψ*** _***c***_ and the arccot curve for ***θ*** of the LOS to e_1_ remains the same. The arccot curve for e_2_ is calculated for an r_2_ = 0.73 cm and skewed by the tct = 1.0 cm. RDUs #1-3 show WF strikes at a-c at e_1_ and d-f at e_2_. (f) 3-D cylindrical polar plot of fig 4(e). Two concentric cylinders with r_1_ and r_2_ radii. Clock hand WF spins around cylinder (green coil). Slices of the cylinder are pulled out to show when the clock hand WF strikes at electrodes e_1_ (blue circles a,b,c) and at e_2_ (red circles d,e,f). The WF strikes at e_1_ are the same is in 4(c), but the WF strike at e_2_ have a much slower progression of increasing time between WF strikes. The diametric property of simultaneous Doppler compression and expansion becomes more asymmetric with skewed electrodes but still exists.

Imaginary lines extending between 3:00 to 9:00 for each of the clocks are parallel lines. It follows then that a clock hand line that extends the center of S to one of the electrodes at the moment of a WF strike, that line is the line of sight (LOS). The times of the clock and the electrode are the exact same time (congruent angles) by the theorem of corresponding angles. The clocks at each electrode are large, such that the clock perimeters hit at the inflection point ***p*** between the 2 electrodes. The radius of the e_1_ clock to its perimeter is the same for the e_2_ clock (r_1_ = r_2_). The vector path of the ***S***_***R***_ is the tangent line to both clocks at e_1_ and e_2_ and intersect each clock’s edge at the same point (**p**) and represents the point of the transverse common tangent (tct) of both electrode clocks in 2D space.

**Table I.**
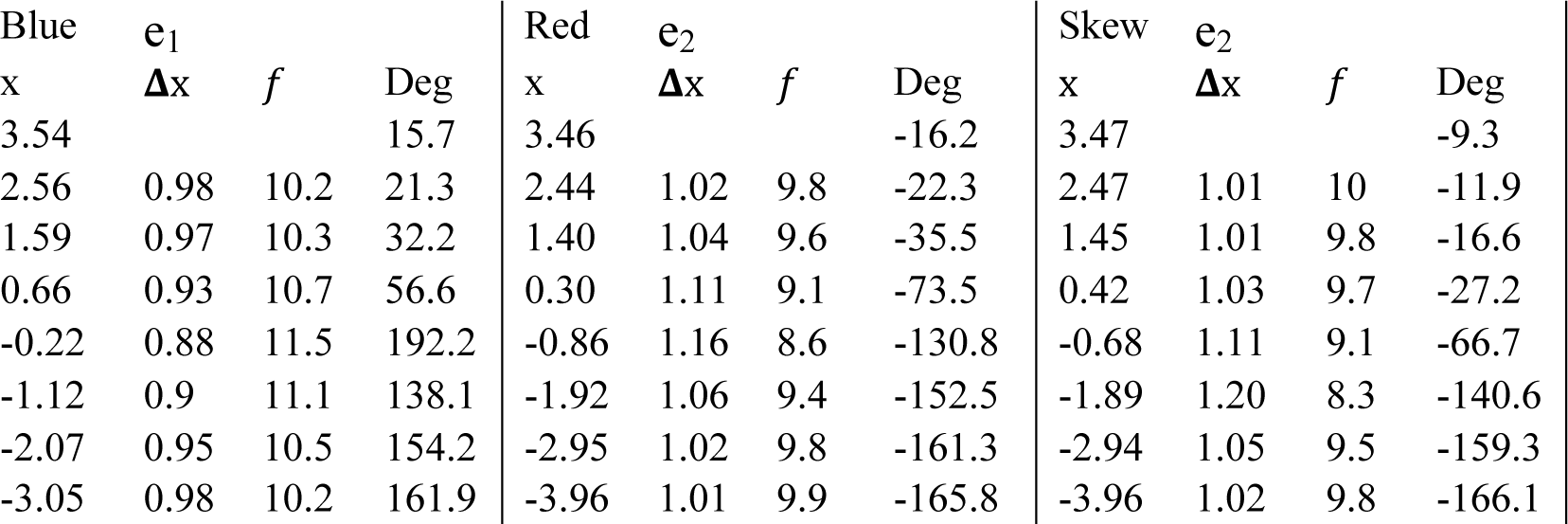
WF strikes positions at e_1_, e_2_ and skewed e_2_ along the ***v***_***s***_ vector path. Values of x represent the distance ***S***_***R***_ is from point ***p*** in Fig 4. Note that as ***S***_***R***_ approaches the inflection point, there is an increasing frequency (*f*) on strong side (blue), while simultaneous decreasing *f* on weak side (red).

| Blue e<sub>1</sub> |  |  |  | Red e<sub>2</sub> |  |  |  | Skew e<sub>2</sub> |  |  |  |
| --- | --- | --- | --- | --- | --- | --- | --- | --- | --- | --- | --- |
| x | $\Delta x$ | $f$ | Deg | x | $\Delta x$ | $f$ | Deg | x | $\Delta x$ | $f$ | Deg |
| 3.54 |  |  | 15.7 | 3.46 |  |  | -16.2 | 3.47 |  |  | -9.3 |
| 2.56 | 0.98 | 10.2 | 21.3 | 2.44 | 1.02 | 9.8 | -22.3 | 2.47 | 1.01 | 10 | -11.9 |
| 1.59 | 0.97 | 10.3 | 32.2 | 1.40 | 1.04 | 9.6 | -35.5 | 1.45 | 1.01 | 9.8 | -16.6 |
| 0.66 | 0.93 | 10.7 | 56.6 | 0.30 | 1.11 | 9.1 | -73.5 | 0.42 | 1.03 | 9.7 | -27.2 |
| -0.22 | 0.88 | 11.5 | 192.2 | -0.86 | 1.16 | 8.6 | -130.8 | -0.68 | 1.11 | 9.1 | -66.7 |
| -1.12 | 0.9 | 11.1 | 138.1 | -1.92 | 1.06 | 9.4 | -152.5 | -1.89 | 1.20 | 8.3 | -140.6 |
| -2.07 | 0.95 | 10.5 | 154.2 | -2.95 | 1.02 | 9.8 | -161.3 | -2.94 | 1.05 | 9.5 | -159.3 |
| -3.05 | 0.98 | 10.2 | 161.9 | -3.96 | 1.01 | 9.9 | -165.8 | -3.96 | 1.02 | 9.8 | -166.1 |

The angle ***θ*** (Figure 3b) is defined as the angle between the LOS and the vector path of the source (vector path arrow points to the zero angle). Therefore, an approaching source is an acute angle until the source reaches the inflection point, (***θ*** at point ***p*** is 90°), at which point the angle ***θ*** becomes obtuse as it recedes from ***p***. The change in ***θ***, or **Δ*θ*** between WF strikes correspond to directly proportional change in the period. Whether one adds or subtracts **Δ*θ*** to the previous position depends upon which side (strong or weak) the electrode sits.

##### b. Determining Strong and Weak Sides: 3-point sequence rule or Wave Front Vector

A sequence-based approach (Fig 3b, right inset) is used to determine if a single observing electrode (e_1_) sits on either the strong or weak side of an approaching ***S***_***R***_. An electrode that sits on the strong side has subsequent shorter periods resulting in higher frequencies of WF strikes. A weak side electrode senses progressively longer periods between WF strikes, resulting in slower frequencies. Two points are placed on the plane of rotation as the initial position of ***S***_***R***_ and e_1_, the spin of S is either CW or CCW. The vector path of ***S***_***R***_ is drawn on plane and divides the plane into 2 sides. Two additional points are specifically placed. One of which is placed directly opposite to e_1_ and is labelled as e_2_. The other point (**e**_**v**_, dashed white circle) is placed on vector path, *forward* of the source’s current position (arrow points to **e**_**v**_, as a reminder). The sequence of the WF strikes over the 3 points is then analyzed. If the sequence of the WF strikes as it sweeps across the 3 points is e_1_-e_v_-e_2_, such that e_1_ always precedes (or leads) **e**_**v**_, then e_1_ sits on the strong side, and e_2_ on weak side. If the sequence was reversed such that e_2_ always preceded e_v_, then the converse is true. Now that one has identified strong and weak sides, one then knows to subtract **Δ*θ*** from the previous period (faster frequency) if electrode on the strong side of an approaching ***S***_***R***_, and adding **Δ*θ*** to the previous period (slower frequency) if electrode sat on the weak side. The 3-point sequence rule is functional whether the spin direction is CW or counterclockwise (CCW) or if the source is approaching or receding from the inflection point.

Alternatively, a vector approach could determine strong and weak sides. If the WF vector was added to the ***S***_***R***_ motion vector (***v***_***s***_), and the sum resulted in an additional positive component in the ***v***_***s***_ direction, then the electrode sat on the strong side. Conversely, if a WF vector added to ***v***_***s***_ resulted in a negative component, then the electrode would be on the weak side. Therefore, subscript labels (st) and (wk) will be used for all subsequent calculations in this paper for side for a moving rotational source. The change in time period **Δ**T can be easily calculated knowing that ***ω***_***s***_ is frequency measurement and the inverse is equal to the period of rotation of the source,

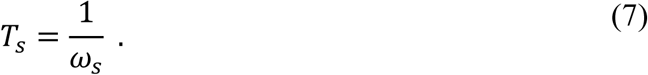

The angular change in the LOS angle ***θ*** between WF strikes results in

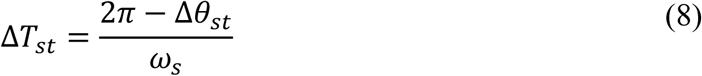

and

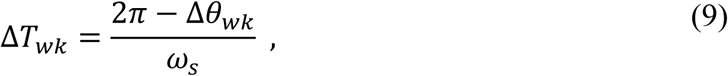

where **Δ*θ*** is a positive value at a strong-side electrode, and negative value at a weak side electrode.

##### c. Rotational math

This method of angular definition and construction provides the proper alignment that when the WFstrikes an electrode, the clock angle ***ψ***_***c***_ is equal to the LOS angle ***θ*** (Fig. 3A and B). Calculating the change that occurs between subsequent WF strikes, that change in clock angles *ψ*_*c*_, allows the direct comparison of the change in time and thus the change in period or frequency between WF strikes.

In the example presented in Figure 3, the center of the emitting source moves at a constant velocity, ***v***_***s***_ = 10 cm/s with a constant ***ω***_***s***_ =10 rotations/s =20*π*/s (in radians). The ***S*** moves along its path that is the perpendicular bisector that approaches and intersects point **p**. The WF from the source has a constant angular velocity. Because ***v***_***s***_ and ***ω***_***s***_ are both constant, during a single rotation (***ψ***_***c***_ ranges from *π* to -*π*), the clock angle changes with distance at a constant rate, ***ω***_s_/***v***_***s***_. Note that time factors out, and that by simplifying the ratio, such that the numerator is 2*π*, then the ratio represents the RDU Eq. 2, and results in

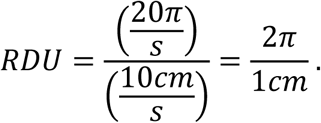

The D value of 1 cm for the RDU shows that one revolution of the source clock is completed with each 1 cm travelled by that source. The RDU can used as the slope of the line that describes how the angle ***ψ***_***c***_ changes as a function of x (the distance from ***p***). This is useful as long as ***v***_***s***_ is not zero. No velocity component would simply result in a vertical slope, the RDU would be undefined, and signify frequency would be constant at ***ω***_**s**_. If one knows the source’s clock position and the source’s clock hand angle, then one could plot the line on a graph (Figures 4A and B) showing the clock hand rotation angle limited from -*π* ≤ ***ψ***_***c***_ ≤+*π* as a function of x (distance from p). At the end of one rotation, when ***ψ***_***c***_ reaches 9:00 or -*π*, then the subsequent rotation would restart at +*π* again as the clock hand continues to rotate at a constant ***ω***_***s***_. This next RDU line would be a parallel, having the same RDU slope from +*π* to -*π*, but the line would be one RDU closer (y-intercept, would advance by 2*π*). This would result in subsequent y-intercepts being advance by 2*π*.

Using the slope-intercept form, for ***ψ***_***c***_ = f(x),

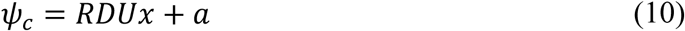

where a represents the y-intercept. The family of RDUs (Fig 4B green lines) with each rotation could be plotted by incrementing the y-intercept by 2*π*, where n represents the integer of the rotation number

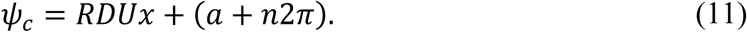

Figure 4 provides a graphical representation of how the ***ψ***_***c***_ as a function the x distance from the inflection point (light green lines Figure 4b). The slope is negative in this CW rotation example with the x-axis reversed a CCW rotation in this graphical presentation would be present as a positive slope. One can use this equation only if the rotation plane of the clock and the electrodes are all in the same plane. Because the source’s vector path is a perpendicular bisector of the line between e_1_ and e_2,_ right triangles can be created between the electrode e_1_ or e_2_, point **p**, and the source location. Superimposed on this graph, the arccotangent curves are plotted which identify the LOS angle ***θ*** along the source’s path of motion to each electrodes e_1_ (blue curve) and e_2_ (red curve). The angle ***θ*** to e_1_ will have a (+) magnitude (blue curve), while ***θ*** to e_2_ will have (-) magnitude (red curve). In this example, an arbitrary starting point for the first RDU, begins with the source clock hand at 9:00 (or *π* radians) with the source 3 cm away from point **p**. The arccotangent function is used instead of tangent since the angle ***θ*** as the source moves past the electrodes will have a range 0 ≤ ***θ*** ≤ *π* for e_1_, and a range of 0 ≥ ***θ*** ≥ -*π* for e_2_. Using the 3-point sequence rule above, e_1_ sits on the strong side of the source’s vector path and e_2_ sits on the weak side such that

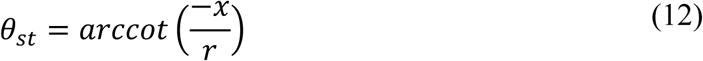

and

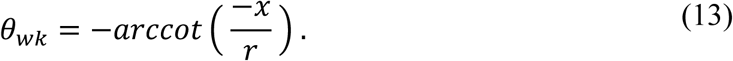

The value of x is made negative to reverse the axis in keeping with a right to left motion of the ***S***_***R***_ towards point ***p***. When one plots ***ψ***_***c***_ and ***θ*** each as a function of ***x*** (Fig 4B), the points of intersection (white circles) identifies when the WF strikes e_1_ and e_2_ where the clock angle ***ψ***_***c***_ is equal to the LOS angle ***θ*** during each RDU.

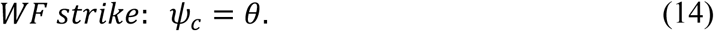

As the ***S***_***R***_ approaches there is one WF strike per rotation (RDU line intersection at arccotangent curve). There is symmetry of the arccot curves about the x-axis. Initially, the timing of a WF strike at e_1_ immediately precedes a WF strike at e_2_. The timing between e_1_ and e_2_ increases with each RDU as it approaches the inflection point (square box, ***θ*** = *π* or -*π*). Once ***S***_***R***_ has passed **p**, the WF strike at e_2_ is so delayed after e_1_, that it now precedes a WF strike at e_1_ of the next RDU line, signaling a reversal in sequence. If one placed e_2_ at positions closer to inflection point (shorter r_2_), then the WF strikes would exhibit less change in time periods at distance far from **p**, but quite more abrupt change with RDUs closest to ***p*** (black curved lines). The change becomes more sudden for closer positioned electrodes to the vector path of the rotational source.

##### d. Alternate method of visualization – cylindrical plot

Creating a 3-dimensional cylindrical polar plot of these functions allows visualization of the clock hand angle to spin around the cylinder, removing separate RDUs, creating a single coil about the cylinder. The arccotan curves that asymptotically approach each other at the 0° at the near end of the cylinder (blue and red solid lines), separate maximally to opposite sides of the cylinder at the inflection point and then approach each other again asymptotically on the other side of the cylinder (blue and red dashed lines). Doppler compression of WF strikes at e_1_ is seen by the subsequent blue circular slices showing the shorter times between strikes. This occurs simultaneous to the Doppler expansion of WF strikes at e_2_ as seen by the successive red circular slices exhibiting progressively longer periods between strikes. The sequence of WF strikes e_1_-e_2_ at the near reverse to e2-e1 once passed the inflection point. This finding provides a trigonometric understanding of the “1/2 cycle drop off” at each moment of phase reversal pattern of WF strikes that we had observed as a cardiac rotor passed through a perimeter of electrodes (*2*).

To gain further insight of this simultaneous diametric property of rotational Doppler effects, an example is shown how the timing WF strikes changes between the same 2 electrodes if ***S***_***R***_ approaches the electrodes from a different angle that does not perpendicularly bisect the line between electrodes (Fig 4(d-f)). The vector path is provided to still have its closest approach to e_1_ be at 1 cm (**r**_**1**_), but is tilted 30° compared to 4(a). A new radius is calculated to its point of closest approach (**r**_**2**_) to e_2_. The distance between the points of closest approach to e_1_ and e_2_ is length of the line segment of the transverse common tangent (tct). The cylindrical transformation can be completed similar to fig 4(b) with two changes. First, constructing 2 concentric cylinders with different radii, and second, skew the inflection points by the tct. The clock hand rotates around through the cylinders intersecting the arccotan curves at the surface of the cylinders (Fig 4(f)), identify the angle of the strike and the distance from the inflection point to that specific electrode.

#### 3. Rotational Doppler effect equation

The diametric property of rotational waves requires the identification of the strong side and weak side of the source’s movement to determine time periods between subsequent WF strikes. Each of the WF strikes from a moving rotational WF source is directly related to subsequent WF angle change between strikes (**Δ*θ***). Strong side changes in the angle of the WF strike results in **Δ*θ*** having a positive value termed now as **Δ*θ***_st_, while the weak side having a negative value, **Δ*θ***_wk_. This allows for the derivation of the rotational Doppler formula.

The change in the clock hand angle ***ψ***_***c***_ at e_1_, between rotation 1 and rotation 2 is **Δ*θ***_st_. Similarly, **Δ*θ***_wk_ represents the clock hand angle change for the WF strikes occurring during rotation 1 and rotation 2 at e_2_. If one knows the constant angular velocity of source’s rotational wave, then the angles **Δ*θ***_**st**_ and **Δ*θ***_**wk**_ equates to the exact time period (T) change between subsequent WF strikes.

The period (***T***) for a rotational wave is defined as the time required to make 1 complete revolution, and is inversely proportional to the angular velocity of the source’s WF,

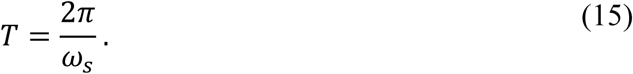

Thus, the angular velocity is frequency measurement is

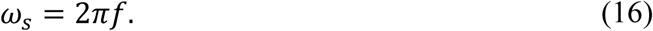

On the strong side of the source’s movement, the subsequent WF strike occurs with less of a complete rotation (less arc length) or 2*π*-**Δ*θ***_st_. The subsequent rotation results in a new period ***T***_***st***_ that is reduced by **Δ*θ***_st_ for e_1_, while ***T***_***wk***_ is increased by **Δ*θ***_**wk**_ or as shown above, such that

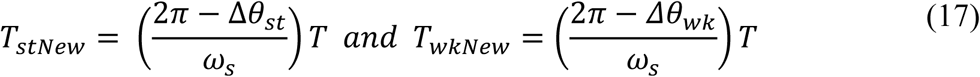

for e_1_ and e_2_ respectively. One can see that subtracting **Δ*θ***_st_ or **Δ*θ***_wk_ (**Δ*θ***_wk_ has a negative value) result in a shorter or longer period between WF strikes respectively. Note that period change with subsequent RDUs, ***T***_***st***_ and ***T***_***wk***_ for e_1_ and e_2_ respectively, are not equal and opposite in time since the WF strikes are not occurring simultaneously at e_1_ and e_2_. The source had moved closer to e_2_ at the time of the WF strike that is recorded there. One can convert the change in period to a change in frequency observed at each electrode since period and frequency are inversely proportional,

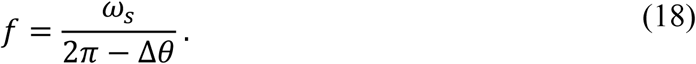

Importantly, **Δ*θ*** is either **Δ*θ***_st_ or **Δ*θ***_wk_, they are not equal, and they are opposite in sign. The new frequency ***f*** ′ can be compared to the original frequency ***f*** ^***º***^, where the derived **Rotational Doppler Equation** (RDE) is now

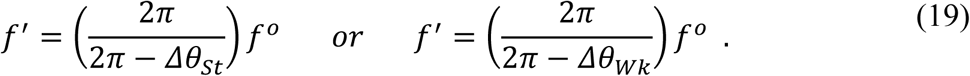

And since ***θ*** = ***ψ*** _***c***_ at WF strikes, the change in the clock hand angle results in

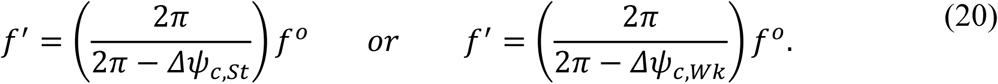

The change in frequency for rotational waves is markedly different for standard doppler such as with sound waves

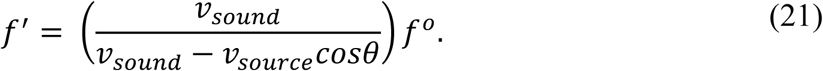

A detailed comparison is made in Fig 5 for different electrode separation from point ***p***, keeping e_1_ and e_2_ equidistant to ***S***_***R***_. Such analysis allows further important general characteristics of the diametric property of a rotational source and can be compared to the diametric property of a spiral source (***S***_***Sp***_) described later.

**Figure 5.**
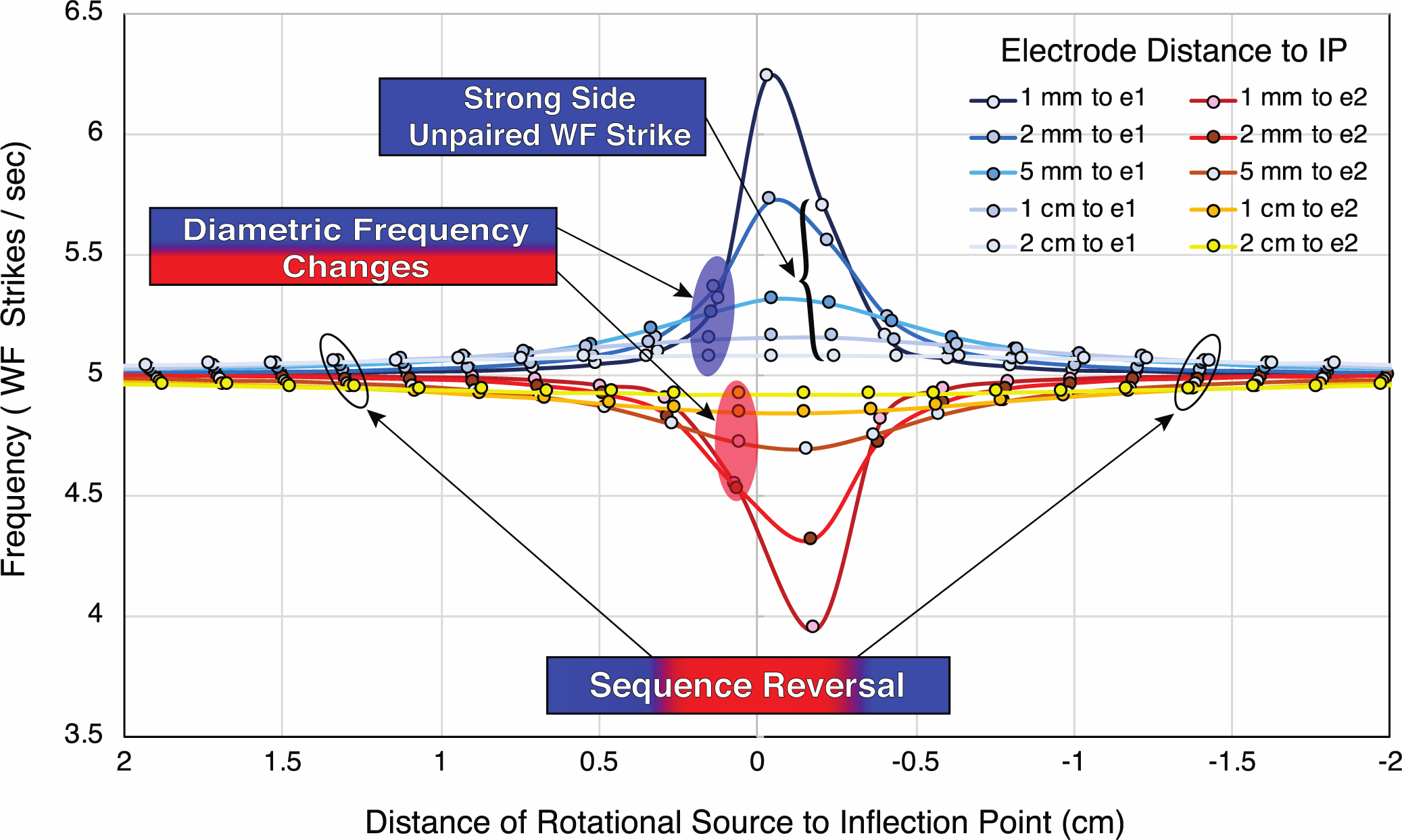
The diametric property of a *S*_*R*_ at different distances of equidistant electrodes. A ***S***_***R***_ has a ***ω***_***s***_ = 5Hz, that moves with ***v***_***s***_ = 1cm/s along a vector path is a perpendicular bisector between e_1_ and e_2_. Electrode pairs sit at 1mm, 2mm, 5mm, 1 cm and 2 cm from the inflection point ***p***. Computed WF strikes are plotted as a function of x from the ***p*** with the instantaneous frequency (hollow circles) on the y-axis. RDU is 2*π*/0.2cm. Frequency effects on the strong side and weak side of the ***S***_***R***_ are shown by smoothed blue shaded curves and red/yellow curves respectively for a particular electrode distance. The approaching ***S***_***R***_ shows that the sequence of WF strikes occur at e_1_ immediately precedes e_2._ After ***S***_***R***_ passes inflection ***p*** (at x=0), the sequence of WF strikes reverses, e_2_ strikes precede e_1_ (hollow black ovals). At about 0.5 cm, marked separation of progressive WF strike frequency occurs comparing strong and weak sides (blue and red shaded ovals). At far left and right of the plot, frequency approaches ***ω***_***s***_ at 5 Hz. An unpaired WF strike on the strong side (black bracket) occurs at the inflection point for each distance. Closer electrodes show more abrupt and greater frequency changes than electrodes that straddle the vector path at further distances. Equations and data from these plots are provide in the Supplementary Materials Figure 5 Data File.

Three main characteristics of the diametric property stand out for an approaching, then receding source of rotational WFs.

1. Reversal of sequence activation at the inflection point.
2. The strong side position of electrodes will sense an increased frequency while simultaneously, weak side positions will sense a decreasing frequency during the approach, with opposite changes occurring when S recedes from ***p***.
3. The strong side will have one greater WF strike than the weak side.

### B. The Derivation of a Spiral Doppler Effect

The cardiac rotor however is not a pure rotational wave, it is spiral. The creation of the RDU in the previous rotational WF example lays the cornerstone to solving WF strike frequency at electrodes of a moving spiral wave. The rotor’s spiral shape is described to most nearly follow that of an Archimedean spiral (*28, 29*) with wave curvature and refractory tissue governing the ultimate period of rotation. A ray drawn from the center of the Archimedean spiral intersects each subsequent arm of the spiral at a constant distance of separation. In Fig 6, an ideal Archimedean spiral is shown. Similar to the clock hand spin, the spiral spin will also exhibit a spin and side dependence to observed frequencies. The Diametric Property of frequency changes exists here as well with movement of the spiral.

**Figure 6.**
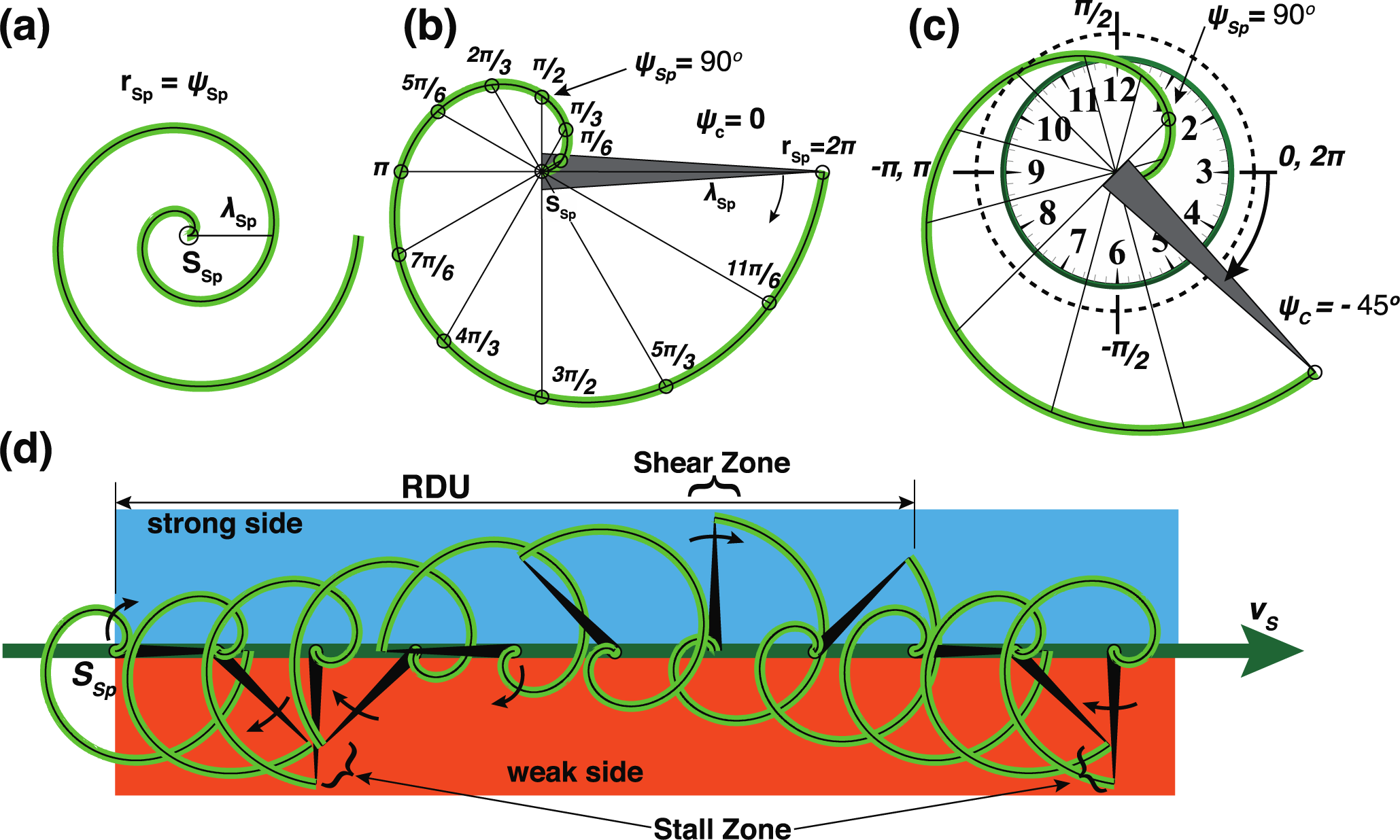
Archimedean spiral and time reference. (a) A source ***S***_***Sp***_ emits a clockwise Archimedean spiral WF governed by equation ***r = ψ*** _**Sp**_ for 2 full rotations with a constant wavelength ***λ*** =2*π*. (b) Expanded view of first revolution of spiral with defining position of clock hand (***ψ*** _***c***_ grey triangle) along the ***λ***-vector line_0→2*π*_. The length of line_0→2*π*_ is the spiral wavelength ***λ*** that defines the size of ***S***_***Sp***_. Radial spokes from center to the spiral arm show the spiral angles ***ψ*** _***Sp***_ at increments of 30°, or *π*/6 radians, and increases in the CCW direction. Clock hand is oriented at the 3:00 position (***ψ***_***c***_ is at zero angle) defined along the spiral source ***S***_***Sp***_ vector path ***v***_***s***_. The point on the WF at spiral angle ***ψ*** _**Sp**_ = 90° is noted at the arrow. (c) As spiral source ***S***_***Sp***_ spins clockwise *π*/4 radians, a point on the spiral wave can still be computed by a spiral summation angle ***ψ*** _***SA***_. The ***ψ*** _***SA***_ is generated by first identifying the clock hand angle ***ψ*** _***c***_ compared to the zero angle and then following the wave back CCW by the spiral angle ***ψ*** _**Sp**_. The clock angle has positive values above the line ***v***_***s***_ and negative values below line ***v***_***s***_. The new position of spiral angle ***ψ*** _**Sp**_ =90° is noted at the arrow. (d) The rotational distance unit (RDU, 2*π*/***D***) is applied to spiral wave. The distance ***D*** travelled is function a constant velocity ***v***_**s**_ of the spiral as it makes one completes revolution at a constant angular velocity, ***ω***_s_. The 3-point sequence rule developed for rotational waves is used here to label the strong side and weak sides of a spiral wave (blue and orange rectangles, respectively). Similar to Fig 3(a), at certain ***ψ*** _***c***_, WF on the strong side would move through a region with faster velocity (shear zones), while on the weak side the WF would move slower (stall zones). Shear zones and stall zones, separated by D, would be expected to repeat all along the vector path on strong and weak sides respectively.

Determining frequency changes of WF strikes from a spiral wave have additional challenges compared to that of rotational waves of Figures 3 and 4. First, the spiral wave, even without spinning, has a changing WF crest angle as one moves away from the emitting source’s center. Second, a spiral wave is one continuous WF crest that has a specific size or wavelength (***λ***). Other than the rotation angle of a cardiac rotor, the description of movement of the spiral WF in a 2-dimensional space is made by the location of its central core as it meanders along the cardiac tissue. For simplicity, it will be assumed that ***S***_***Sp***_ will maintain its spiral shape for at least 2 wavelengths out from its center, or 2 complete turns, as it moves along its vector path ***v***_***s***_. In the cardiac rotor, WF head (crest of WF) meets tail (absolute refractory period) the distance ***λ*** (between 2 arms of the spiral) should not be able to be compressed beyond this physical limit (see Discussion). Experimental studies suggest the cardiac rotor ***λ*** may be in the range of a few hundred microns up to 2-3 cm (*7, 28, 30*). The calculations and plots of ideal spirals with ranges of similar ***λ***s below provide an accurate approximation for typical rotor frequency as they would be recorded from electrodes.

#### 1. Spiral math

One must first develop a spiral math from the general spiral equation

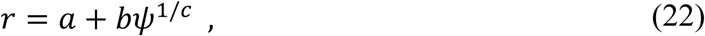

to plot the WF crest as it spins and moves, where radius distance r is the distance from the center to a point on the spiral, ***a*** turns the spiral, ***b*** is the spiral size, and ***ψ*** is the angle of rotation from a zero angle to point on the spiral arm. For Archimedean spirals, c=1, and the polar wave equation is

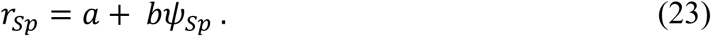

##### a. Creating a spiral clock, ψ _c_

The size of an Archimedean spiral is determined by a measurement from the center of the spiral to the point where the spiral has completed one full rotation (2*π* radians). This distance is the wavelength, ***λ***, and is therefore also the distance between all subsequent spiral arms (Fig 6). Setting ***a*** = 0, a ray that extends from the center of the spiral to a point on the spiral (r = ***λ***) at one revolution (***ψ*** _***Sp***_ = 2*π* radians) then the spiral size is

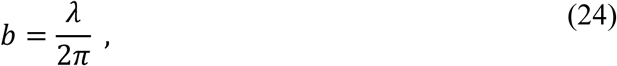

and solving for ***ψ*** _***Sp***_, the equation which reconfigures to

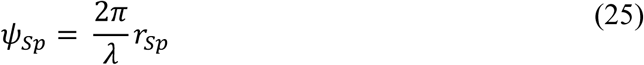

and shows that the spiral angle ***ψ*** _***Sp***_ is inversely proportional to ***λ*** of spirals of different size. If spiral A is twice the size of spiral B, a line segment equal to spiral A’s wavelength (2*π*) would hit the spiral B’s curve at 4*π* or at two full revolutions. A spiral angle *ψ* _***Sp***_ can be calculated for a specific spiral source ***S***_***Sp***_ of size ***λ***, where a ray, ***r*** _***Sp***_ is any fraction or multiple of ***λ*** _***Sp***_.

From here forward, ***S***_***Sp***_ will identify a source that emits a spinning Archimedean spiral wave governed by the Eq. **19.** Movement of the spiral will be defined by 2 velocities, the velocity of the center of the spiral along the vector path (***v***_***s***_), and the angular velocity of its spin (***ω***_**s)**_. Two angles, the clock angle ***ψ*** _***c***_ and the spiral angle ***ψ*** _***Sp***_, must be determined to make frequency calculations of WF strikes from a spiral wave. The clock angle ***ψ*** _***c***_ identifies the specific position in its rotation that is referenced to a zero angle in 2-D space, while the spiral angle ***ψ*** _***Sp***_ identifies the local angle of a specific position on the spiral wave referenced to its immediate spin position ***ψ*** _***c***_. As seen in Fig 6, a spiral that spins in a clockwise direction, its spiral angle ***ψ*** _***Sp***_ of Eq. 2 increases in the opposite, or counterclockwise direction as one measures progressively further from the spiral center along the spiral arm. Conversely, a counterclockwise spinning spiral will have a spiral angle ***ψ*** _***Sp***_ that increases in a clockwise direction.

An Archimedean spiral replaces the clock example of Figure 3 and 4 above. A line segment_0→2*π*_ can be drawn on the spiral from its center to a point on the wave where it has completed one full rotation (r = 2*π*). An imaginary clock hand is glued along this segment (line_0→2*π*_ grey triangle, 6(a) and (b)). The length of the line segment from r = 0 to r = 2*π* is equal to the wavelength ***λ***_***Sp***_. A ***λ***-vector line describes the direction of line_0→2*π*_ in 2-D space. As in the prior rotational example (Fig 3A), the ***S*** _***Sp***_ vector path ***v***_***s***_ points to the clock’s 3:00 position and is used as a referenced zero angle. Therefore, the clock hand angle ***ψ*** _***c***_ for a spiral wave will be defined as the angle between the ***λ***-vector line and vector path ***v***_***s***_ (Fig 6(b) and (c)).

##### b. Spiral summation angle, ψ _SA_

Any point on the spiral wave can now be described by a spiral summation angle (***ψ*** _***SA***_) of ***ψ***_***c***_ and ***ψ*** _***Sp***,_

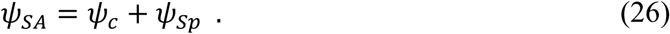

In this example, spiral ***S***_***Sp***_ spins at a constant angular velocity, ***ω***_s_, that turns the clock hand angle ***ψ*** _***c***_. Any stationary observer that is placed on this 2-D plane will have its clock perfectly synchronized to that of ***S***_***Sp***_. The change in the spiral’s clock angle ***ψ*** _***c***_ between any two sequential WF strikes of the spiral arm at the observer is a measure of the time period (**T**) between WF strikes and can be converted to a frequency measurement. The **T** between WF strikes will be measured by the spiral’s clock angle ***ψ*** _***c***_, since the WF crest is a changing curve at the observer. If the spiral spins, but its center is stationary (***v***_***s***_ = 0), the WF from ***S***_***Sp***_ will strike the stationary observer with a constant **T**, no matter where it is positioned with each subsequent rotation. The frequency of WF strikes would be constant and equal to ***ω***_s_. With each one revolution, the WF crest moves one ***λ*** away in distance over time **T** out at a constant rate. If that same spinning ***S***_***Sp***_ center is put in motion at a constant ***v***_***s***_ (along this x-axis), then the distance ***S***_***Sp***_ travelled, **D** (or ***v***_***s***_ T of the RDU, Eq. 5) is also a measure of **T** along the x-axis during that one revolution. Since both ***ω***_***s***_ and ***v***_***s***_ are constant, the changes in **T** between WF strike at an electrode can be calculated from either the change in angle ***ψ*** _***c***_ at the spiral or the change in the x-distance that the spiral moved between WF strikes. Using distance as a measure of time is the source’s distance-time perspective. The spiral’s time measurements for calculation of frequency will be the same as that perceived by the electrodes. In addition to electrode distance from a spiral’s path, the wavelength of the spiral can also be varied to assess the diametric property.

The ***v***_***s***_ vector path of ***S***_***Sp***_ in Fig 6(d) separates 2-D space in half. The 3-point sequence rule still applies here to designate specific sides. If 2 observers were placed one on either side of ***v***_***s***_, then the strong side and weak side nomenclature holds for an approaching ***S***_***Sp***_. WFs from the spiral move faster through the strong side and slower on the weak side. On the strong side, at a certain radial distance from the spiral center, a spiral arm may propagate at its fastest forward velocity allowed by the substrate. As the spiral arm turns the WF may shear from its outer arm, developing as a shear zone on the substrate. Conversely, on the weak side, at certain radial distance from the spiral center, the WF may appear to linger and pivot, manifesting as a stall zone on the substrate. These repeating regions of shear and stall, within the strong and weak side respectively, are staggered along the ***v***_s_ path separated by the distance D of the RDU. Examples of likely phenomena of shear and stall zones will be discussed below.

Similar to the example presented in Fig 3 and 4, and to prove the diametric property exists for spiral waves as well, stationary electrodes (e_1_ and e_2_) are placed such that the vector path ***v***_**s**_ of ***S***_***Sp***_ is a perpendicular bisector between them (Fig 7). Electrodes e_1_ and e_2_ are equidistant to the center of the spiral ***S***_***Sp***_ all along its path. 2 observers (electrodes). ***S***_***Sp***_ spins with a constant angular velocity ***ω***_s_. The activation sequence of WF strikes at electrodes e_1_ and e_2_ that straddle either side of ***v***_***s***_, must reverse their sequence as the spinning ***S***_***Sp***_ passes the inflection point (***p***) of the transverse common tangent. The LOS angle, ***θ***, remains the angle between a line from the center of ***S***_***Sp***_ to e_1_ or e_2_ and the line of the forward vector path ***v***_***s***_. The LOS angle ***θ*** ranges from 0° to 180° as the source moves from being an approaching source to a receding source from an inflection point. LOS angle ***θ*** is positive if the electrode is above the vector line ***v***_***s***_, and negative if the electrode sits below ***v***_***s***_. Each electrode sits at the center of imaginary perfectly circular clocks created in the same 2-D plane (blue and red circles). The clock perimeters are of specific radial size, such that ***v***_***s***_ intersects the transverse common tangent to both clock’s perimeter at point ***p***. Point ***p*** is therefore a common inflection point (***p***) where the moving ***S***_***Sp***_ changes from being an approaching ***S***_***Sp***_ to a receding ***S***_***Sp***_. The measured distance ***h*** along the LOS angle ***θ*** to either e_1_ or e_2_, is simply the hypotenuse of a right triangle that is created from ***S***_***Sp***_, ***p***, and either e_1_ or e_2_,

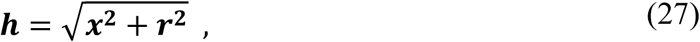

where r is the radius of the imaginary clock perimeter of e_1_ or e_2_, and ***x*** is the distance from the ***S***_***Sp***_ to the inflection point **p**. A positive value of ***x*** is used as the ***S***_***Sp***_ approaching ***p*** and negative if receding from ***p***. Although the value of ***x*** is squared here, it’s positive and negative values will need to be remembered and used later with frequency calculations. Since e_1_ and e_2_ are equidistant to the spiral S all along its path, the LOS angle ***θ***, is also equal all along its path. Any diametric changes (increasing vs decreasing) in frequency detected by the electrodes will be a result of strong and weak side of spin direction of the spiral.

**Figure 7.**
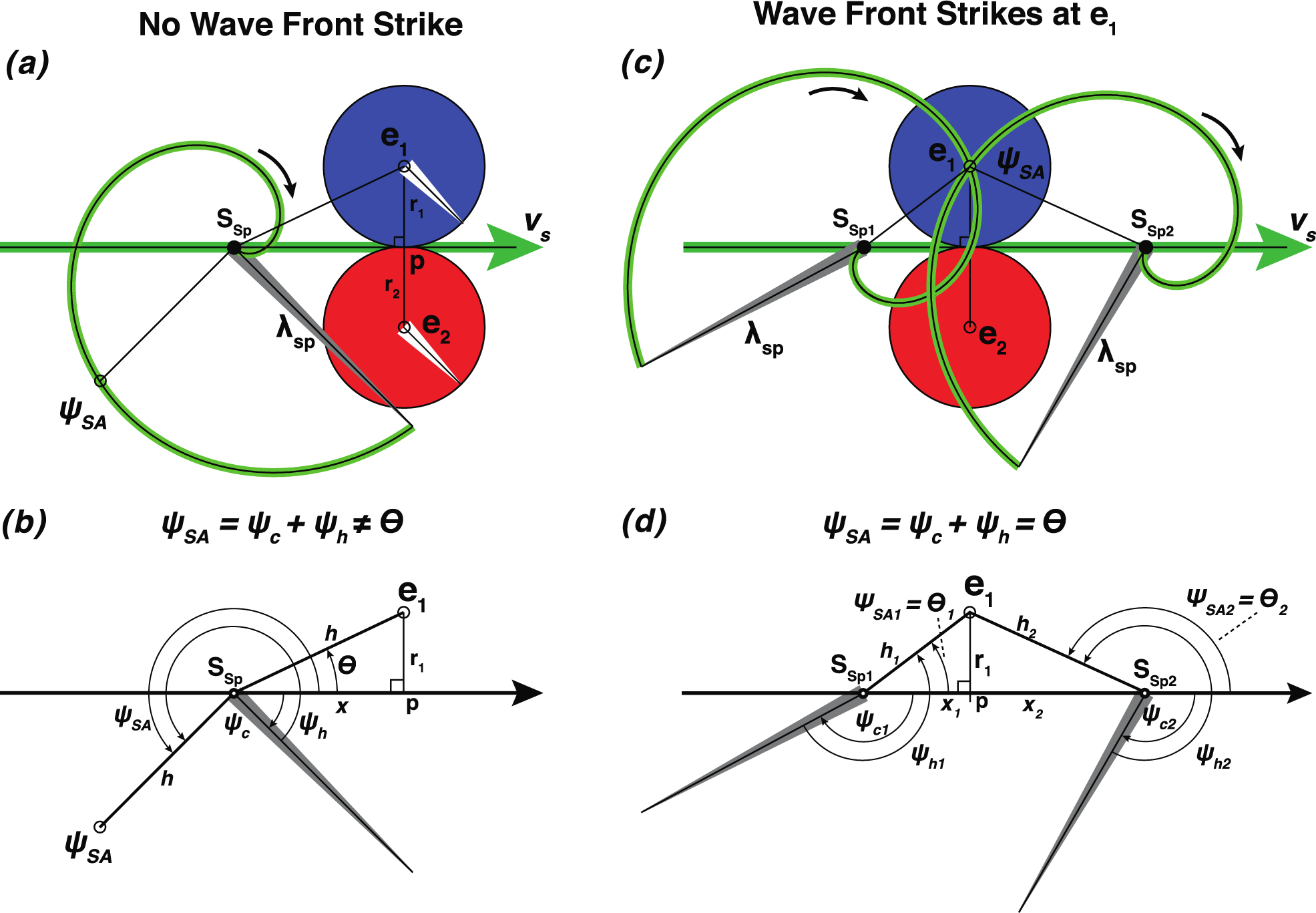
Spiral math components determine WF strikes. (a) and (b). A snapshot of a moment in time that a clockwise spinning spiral source ***S***_***Sp***_ approaches the inflection point (***p***) between 2 electrodes e_1_ and e_2_ (center of blue and red circles, respectively) that are equally spaced across ***v***_***s***_ (r_1_ = r_2_), on either side of the vector path ***v***_***s***_ of the source. The Archimedean spiral WF crest (green striped curve) completes one full turn, with a wavelength ***λ*** _***Sp***_ = 2*π*. **Step 1**: determine the clock hand orientation (thin gray triangle) of the spiral by the ***λ***-vector line on the ray from the center of the spiral to the point of one complete revolution. The clock hand angle, ***ψ*** _**c**_ is measured by placing the zero angle along the forward vector path, ***v***_***s***_ (same as in Fig 3A). **Step 2**: determine the distance from S to e_1_, the hypotenuse ***h***, of the right triangle S-p-e_1_. The distance ***h*** is converted to the angle ***ψ*** _**h**_ and is plotted on the spiral curve in a counterclockwise direction from ***ψ*** _**c**_. The resultant point on the spiral, at the ***ψ*** _**SA**_, in 2-D space identifies that the spiral WF crest did not intersect with e_1_ at that position and time. (c) and (d). Two snapshot examples (approaching and receding) are shown where the spiral WF crest intersects with e_1_, where the source at position ***S***_***Sp1***_ approaches the inflection point ***p*** and where the source at position ***S***_***Sp2***_ recedes from point ***p***. At ***S***_***Sp1***_, the clock hand angle ***ψ*** _**c1**_ is added to the hypotenuse angle ***ψ*** _**h1**_ resulting in an acute angle ***ψ*** _**SA**_ that is equal the acute LOS angle ***θ***_**1**_. At ***S***_***Sp2***_, the clock hand angle ***ψ*** _**c2**_ is added to the hypotenuse angle ***ψ*** _**h2**_ resulting in an obtuse angle ***ψ*** _***SA***_ that is equal the obtuse LOS angle ***θ***_**2**_. Close inspection of ***S***_***Sp1***_ shows that the WF strike at e_1_ has occurred before a WF strike would occur at e_2_, whereas the sequence is reversed when source is passed ***p*** at position ***S***_***Sp2***_. Here, the WF strike at e_2_ had already occurred just before the strike at e1.

##### c. Hypotenuse used as a spiral angle, ψ_h_

To determine when the WF strike occurs of an Archimedean spiral at an electrode, the size of the spiral, spin orientation, position along its vector path must be known. As the source ***S***_***Sp***_ moves closer to inflection point ***p***, the distance to the electrode ***h*** (from Eq. 24 above), can be substituted into the equation for ***r***_***Sp***_ to determine exactly where on the spiral the translated hypotenuse angle, denoted as *ψ*_***h***_ exists,

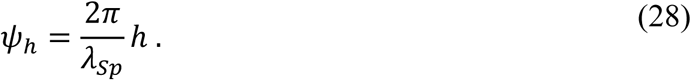

One can add the hypotenuse angle *ψ*_***h***_ at that moment in time to the clock angle *ψ*_***c***_ for that specific ***x*** position of ***S***_***Sp***_. That sum of the angles, a spiral summation angle *ψ* _***SA***_ is

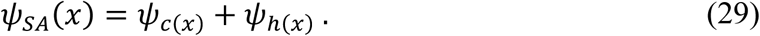

The *ψ*_***SA***_ can then be compared to the LOS angle ***θ*** (Fig 7(a) and (b)). One can see that only when the *ψ* _***SA***_ equals the LOS angle ***θ*** will the spiral WF strike the electrode (Fig 7c and d). As described above, the change in *ψ*_***c***_ between WF strikes at the first position (***S***_***Sp1***_) and the second position (***S***_***Sp2***_) or a measure of the distance x travelled by ***S***_***Sp***_ can be computed as a time period or instantaneous frequency. As seen in Figure 7c, there is a sequence reversal of WF strikes at e_1_ and e_2_ from a position ***S***_***Sp1***_ that approaches ***p*** to a position ***S***_***Sp2***_ that recedes from point ***p***. A sequence reversal indicates that the frequency changes at e_1_ must be opposite to frequency changes at e_2_ and therefore the diametric property of strong and weak sides must also exist for spiral waves.

##### d. Movement of spinning spiral wave and plotting spiral summation curves

To analyze the diametric property changes in WF strike frequency in detail, the snapshot examples of Fig 7 needs to be advanced to motion of the spiral as a function of x as the spiral both spins and moves. For spiral waves, one plots both the family of ***ψ*** _***SA***_ for each rotation (similar to family of RDU in Fig 4) and the LOS angle ***θ*** as a function of ***x*** (example provided in Fig 8). Each clock angle ***ψ***_***c***_ as a function of x is the same calculation of RDU lines as Eq. 2 reduced to the numerator equaling 2*π*. The plot is again limited from -*π* ≤ ***ψ***_***c***_ ≤+*π* such that at the end of one rotation, when ***ψ***_***c***_ reaches 9:00 or -*π*, the y-intercept and is incremented by 2*π*. The hypotenuse angle ***ψ*** _***h***_, converted from the ***h*** distance of ***S***_***Sp***_, is added to the clock angle ***ψ*** _***c***_ for all values of ***x***. Substitution of the component parameters of ***ψ*** _**c**_ and ***ψ*** _**h**_ into Eq. 26

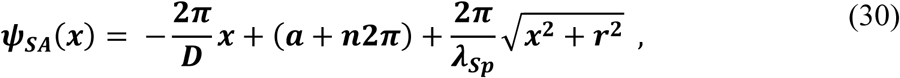

which can accurately identify where on the spiral WF crest a distance ***h*** is measured from the spiral wave’s center at any specific x-position and spin orientation. The y-intercept component essentially sets the clock time at a specific portion of an RDU. Every multiple ***n*** distance D from that position on the x-axis will have the same ***ψ***_***c***_. The value ***a*** is within the range 0≤ *a* ≤ 2*π*, and ***n*** is an integer from -∞ < n < +∞. For one specific ***ψ***_***SA***_ curve, one could replace that y-intercept with the variable K for a where K = a +n2*π*, and simplifies equation 30 to

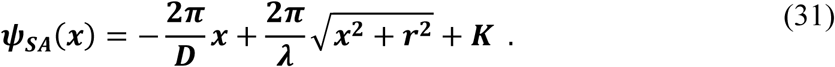

**Figure 8.**
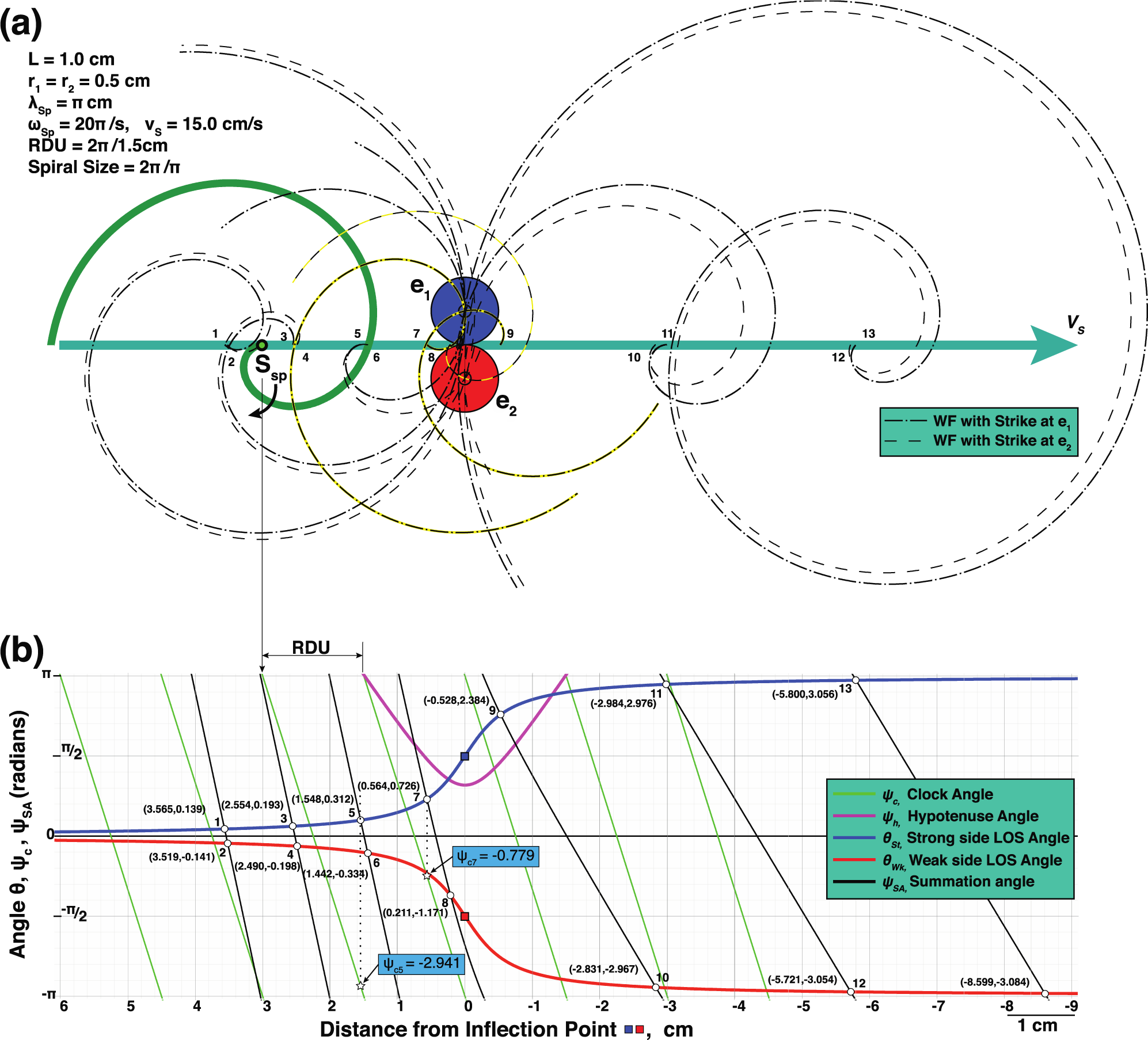
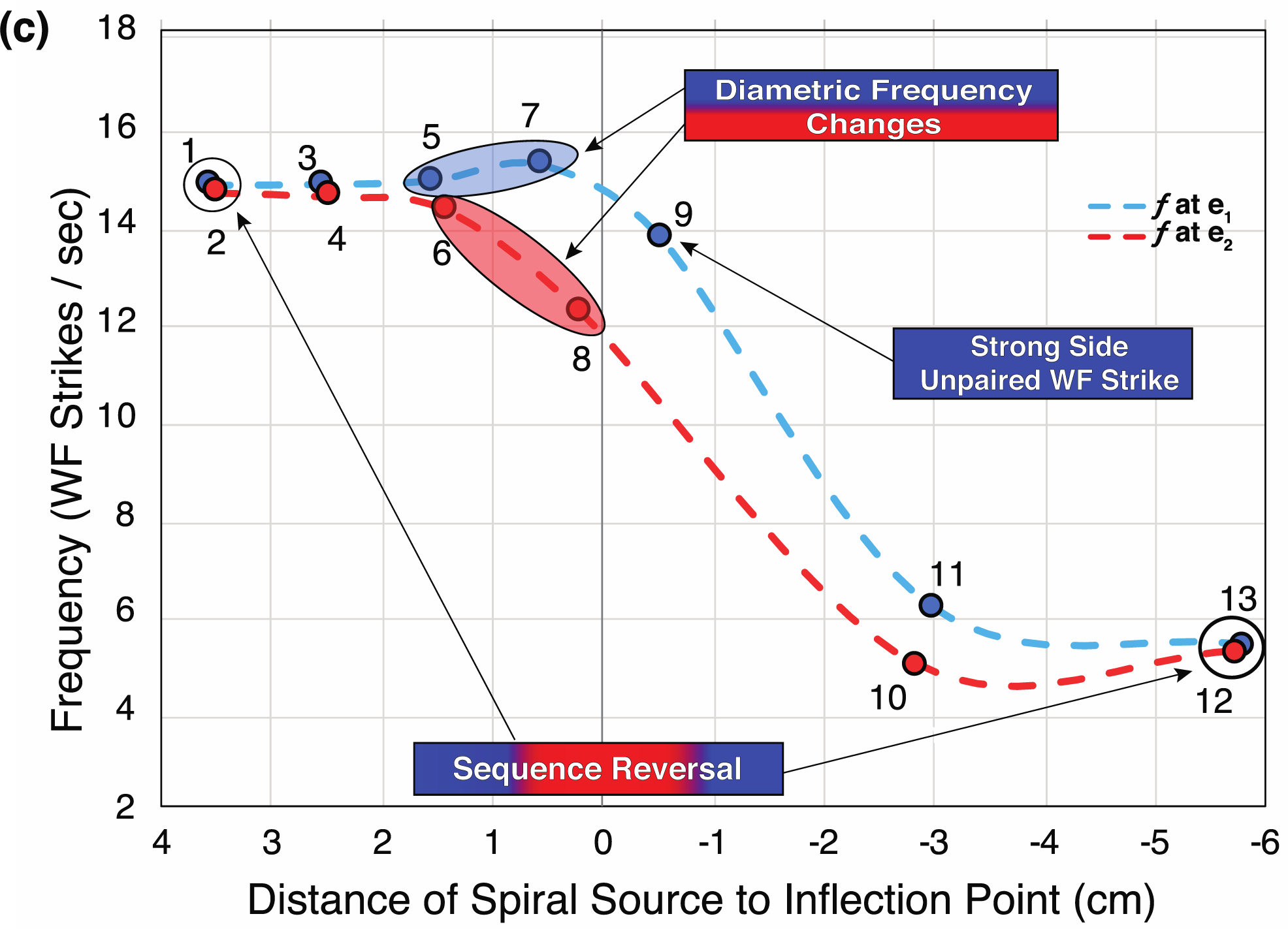
Equidistant electrodes on either side of source’s vector path reveals the diametric property of spiral WF Doppler effects. ***(a) S***_***Sp***_ has a wavelength of *π* radians, a spiral size ***λ*** = *π*cm and spins with an angular velocity of 10 rotations/s (20*π*/s). ***S***_***Sp***_ moves on a vector path at *v*_*s*_ = 15 cm/s that is a perpendicular bisector between electrodes e_1_ and e_2_, each 0.5 cm from the point of closest approach (transverse common tangent to blue and red circles). A reference position of ***S***_***Sp***_ at 3 cm (green spiral) from the inflection point between electrodes is the spiral source’s clock angle ***ψ*** _***c***_ is at 9:00 or *π* radians. The spiral wave in its x position and spin orientation is shown at each WF strike as calculated to occur at positions 1-13. WF strikes at e_1_ (bold dash-dot spirals) are compared to WF strikes at e_2_ (dashed spirals). Black and yellow dashed spirals show the sequence reversal of WF strikes secondary to the diametric property by opposite and progressive changes of **T** comparing strong and weak sides (positions 7-9). Strong side has one more WF strike than weak side (unpaired WF strike at position 9), identical to that seen during rotor migration breaching perimeter of electrodes (*2*) that we described as a ½ cycle drop-off. See Supplementary Materials Figure 8 Data File. (b) The LOS angle ***θ*** is plotted as a function of distance x to inflection point by the arccotangent curves for electrodes e_1_ and e_2_ (blue and red curve, respectively). Superimposed on graph is the spiral clock angle *ψ*_*c*_ (green lines), the distance to electrodes in radians as the hypotenuse angle ***ψ*** _**h**_ (small portion of the pink parabolic curve is illustrated), the spiral summation angle ***ψ*** _**SA**_ (black lines). The points of intersection of ***ψ*** _**SA**_ with the arccotangent curves (white circles) indicate when ***ψ*** _***SA***_ = ***θ***, identifying a WF strike. When ***S***_***Sp***_, approaches inflection point, times between WF strikes for both e_1_ (***S***_***Sp***_ positions at 1,3,5,7) and e_2_ (***S***_***Sp***_ positions at 2,4,6,8) are shorter (higher frequency) than the times between WF strikes when ***S***_***Sp***_ recedes from inflection point for both e1 (***S***_***Sp***_ positions at 9,11,13) and e_2_ (***S***_***Sp***_ positions at 9,11,13). Example for frequency determination at WF strikes at e_1_ from position 5 to position 7 can be calculated by either **Δ*ψ*** _***c***_ or **Δ*x***. The **Δ*ψ*** _***c***_ method would be via using ***ψ***_***c***_ values at hollow stars (−2.941, -0.779, respectively) and plugging into frequency Eq. 35, or by the **Δ**x method by equation 36. (c) Side-Dependent Frequency Changes of Spiral Wave. Plots of the instantaneous frequencies at each WF strike are shown for both strong and weak electrodes (e_1_ and e_2_, respectively). The approaching spiral wave shows a progressive increase in frequency of WF strikes at the strong side electrode e_1_. Simultaneously, there is a progressive decrease in frequency of WF strikes at the weak side electrode e_2_. This diametric progression leads to an unpaired WF at e_1_ (position 9) resulting in the sequence reversal with WF strikes at e_2_ preceding e_1_. The frequency of WF strikes by an approaching ***S***_***Sp***_ must change differently between strong and weak sides as the sequence must reverse once past the inflection point. From relatively far distances (x>>r), strong and weak sides frequencies of WF strikes be nearly the same, at 14.78 WF strikes /sec. When ***S***_***Sp***_ recedes from ***p***, frequencies of both strong and weak sides then approach a final receding frequency of spiral 5.22 WF strikes/sec as per Eq. 40 or 10 ± 4.78 WF strikes/sec.

Thus, each spiral summation curve is made up of two components for its slope, clock spin orientation RDU and spiral size-specific distance.

#### 2. Spiral Doppler effect Equation

In this first example provided in Fig 8, the parameters of ***v***_***s***_ and ***ω***_***s***_ were chosen to allow separation of the sequential WF strikes along a vector path to more easily visualize the diametric property. Each spiral curve plotted (Figure 8a) are at each moment that a WF strikes occur at e_1_and e_2_ (bold dot-dash spirals, thin dash spirals respectively). The stationary positions of e_1_and e_2_ are such that they are equidistant to the center of the spiral all along its path of motion. The points of intersection (white dots, Figure 8b) between the family of ***ψ*** _***SA***_ lines (black lines) with the arccotangent curves for the LOS angle ***θ*** to electrodes e_1_and e_2_ (Eq. 9 and Eq. 10, blue and red lines respectively) indicate exactly when a WF strike occurs (***ψ*** _***SA***_ = LOS ***θ***). The distances (and time periods) between WF strikes at both e_1_ and e_2_ prior to inflection points (blue and red squares), are comparably shorter than the distances between WF strikes once past the inflection points, as reflected by the calculated frequency plots in Figure 8c. More important than comparing an approaching and receding position of the spiral wave, is the comparison of which side (strong or weak) the electrode sits (Figure 8c). As the ***S***_***Sp***_ approaches both electrodes of same distance of separation, electrode e_1_ increases in frequency simultaneously to a decrease in frequency at e_2_.

As a point of reference, the family of RDU lines or ***ψ*** _***c***_ (green solid lines) are plotted. Only a portion of ***ψ*** _***h***_ is plotted to minimize plotted lines.

These initial motion parameters were chosen to provide less WF strikes and simplify the visualization of the diametric property. WF strikes occurred with each rotation when ***ψ*** _***SA***_ = ***θ***. The clock angle component of ***ψ*** _***SA***_ provides the solution to frequency and must be differentiated from the clock angle of a straight arm rotational WF. The clock angle ***ψ***_***c***_, for WF strike calculations of the RDE is a direct function of ***θ***,

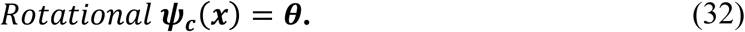

To avoid confusion, by adding a superscript Sp to ***ψ***_***c***_, the distinction can be made where WF strikes calculation to assess the spiral Doppler effect, is a function of ***ψ*** _***SA***_ – ***ψ*** _***h***_,

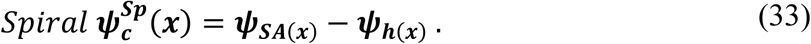

The RDE equation can now be modified to a spiral solution,

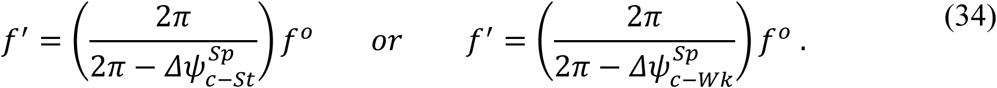

More simply the **Spiral Doppler Equation (SDE)** is,

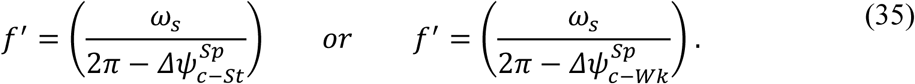

The frequency calculations from WF strikes can also be calculated most easily by using the spiral ***λ*** distance-time, D as a scale of frequency, and the change in x-values between WF strikes measured against D. An **alternate SDE** is,

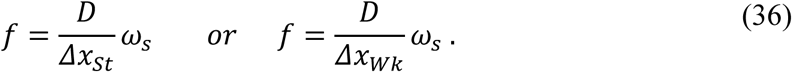

##### 1. Special case of electrode sits directly on path of S_Sp_

Two changes occur with frequency calculations when an electrode sits directly on the ***v***_***s***_ vector path of ***S***_***Sp***_. First, the WF strikes would occur with LOS ***θ*** = 0, no matter the distance from inflection point ***p***. The subsequent WF strikes, in this special case, would occur only when ***ψ***_***SA***_ = 0 radians. ***ψ***_***SA***_ is independent of the distance from ***p***, and therefore not on a changing ***θ*** governed by a side-specific arccotangent curve. Second, r=0, within the spiral size-specific distance component of the Eq. 31. The ***ψ***_***SA***_ equation simplifies to

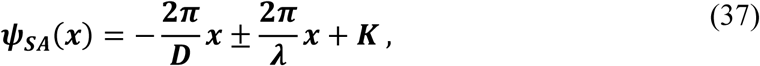

where the (-) sign is used as ***S***_***Sp***_ approaches the electrode, and (+) as ***S***_***Sp***_ recedes from it. The solution results in 2 families of parallel straight lines, approaching and receding. The slopes of the line or first derivative provide the ability to make a calculation of frequency,

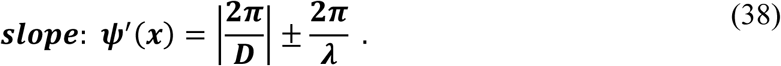

An absolute value of the D component of the RDU is now used to convert the reversed x-axis that was originally needed to measure clock angles ***ψ***_***c***_ in Fig 3, 4, 7, and 8. The slope on the polar graph can be converted to a frequency by multiplying the slope by ***v***_***s***_/2*π*,

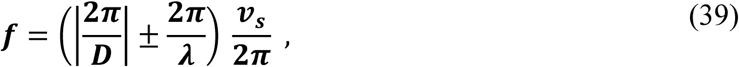

or

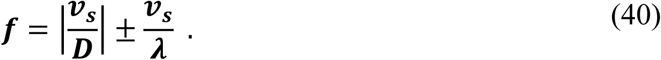

This equation also provides the same result in frequency for electrodes not on the vector path ***v***_**s**_ when ***S***_***Sp***_ is far away from the inflection point. The ***r*** value becomes insignificant in the calculation of the hypotenuse angle ***ψ*** _***h***_ of Eq. 31. The WF strikes, intersection points on the arccotangent curves approach asymptotically the x-axis (approach y = 0). It is only at relatively short distances of ***S***_***Sp***_ to the inflection point, that frequency changes, the diametric property would be observed.

Both components of slope of Eq. 38 have numerators of 2*π* which allows direct summation of angles in a polar perspective. It will be shown below that by constructing this equation in this manner, D and ***λ*** become an important ratio with regard to frequency of WF strikes when comparing approaching and receding spirals.

## III. RESULTS

### A. Electrode separation effect on diametric property of SDE

Cardiac rotors during atrial fibrillation move about tenfold slower than in the teaching example of Fig 8 above. Typical human cardiac rotors that have angular frequencies ranging from 4-8 Hz (*31*), and can drift at 10-20 mm/sec (*7*). The diametric property is more apparent at these slower ***v***_**s**_ rates. Computed comparisons of various electrode separation and spiral sizes are shown in Fig 9 and Fig 10 below to help identify more specific characteristics of the diametric property of the spiral Doppler effect. In both figures, a spiral WF source, ***S***_***Sp***_ moves at a velocity of 1 cm/sec between 2 equidistant electrodes with an angular velocity of 5 Hz. If an electrode sat directly on the ***v***_***s***_ vector path, the approaching and receding WF strike frequency would be 5.32Hz (T=188 msec) and 4.68 Hz (T= 213 msec), respectively (using Eq. 40). Simultaneously, just 2 mm to the side of the vector path (Fig 9 dark blue and dark red curves), the diametric property of the SDE shows that the strong side has a peak WF frequency of 5.78 Hz (T= 173 msec) while the weak side has a nadir frequency of 4.24 Hz (T= 238 msec). Electrodes positioned closest but to the side of the vector path of a spinning and moving ***S***_***Sp***_ have the greatest magnitude changes in frequency than electrodes spaced further out or if they had been placed directly in front or back. For a moving source, ***S***_***c***_, that emits a periodic WF in a classical outward centrifugal propagation, then the electrodes spaced equidistantly on either side of the vector path ***v***_**s**_ would experience WF strikes simultaneously and the change in frequency would be governed by the classical Doppler Eq. 21 as seen in Fig 9B. Peak and nadir frequency are seen directly in front or behind, not to the sides as that of a spiral source.

**Figure 9.**
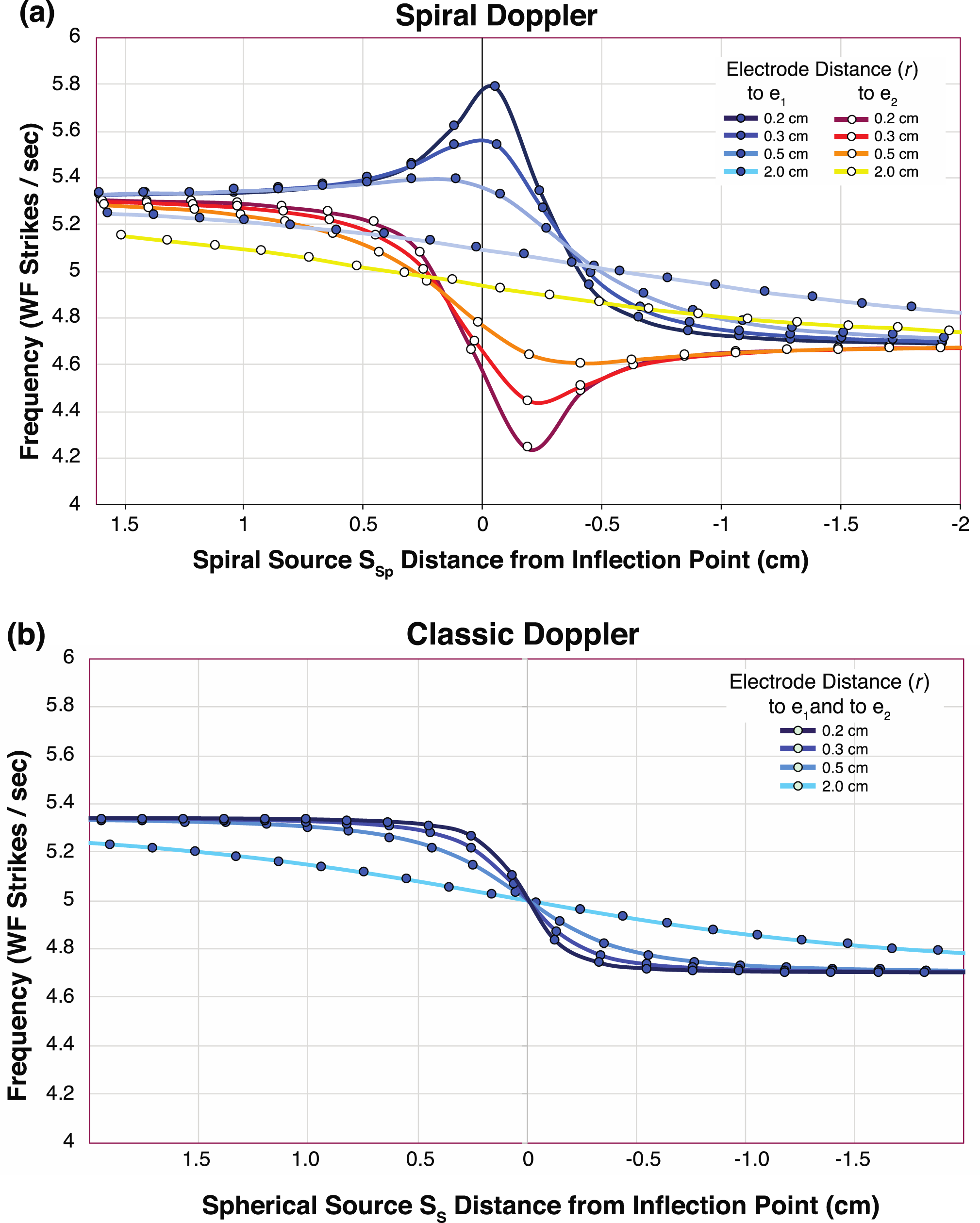
Comparison of frequencies observed from a moving spinning spiral WF source and a moving source that emits centrifugal propagating WFs. (a). Comparison of frequency changes with propagation of a spiral wave between different distances of equidistant electrodes. The speed of spiral source ***v***_***s***_ =1 cm/sec. Angular frequency is 5 rotations/sec, RDU = 0.2 cm^-1^, with ***λ*** = *π*cm. Comparison of calculations were made with stationary electrodes placed at 0.2 cm, 03 cm, 0.5 cm, and 2 cm from the inflection point ***p***. The closer the electrodes were to the ***S***_***Sp***_ center, the more abrupt the change in *f* and the larger the difference between the strong and weak side electrodes (gradation of darker blue and darker red lines, respectively). Closest electrodes show an increasing *f* on the strong side, simultaneous to a decreasing *f* on the weak side of ***S***_***Sp***_. At larger electrode separation distances, the approaching distant ***S***_***Sp***_ has a *f* faster than ***ω***_***s***_ that gradually decreases, and becomes less than ***ω***_***s***_ after receding from ***p*** (light blue and light yellow lines, respectively). From a distance the using Eq. 40, the approaching *f* = 5.32 Hz and the receding *f*= 4.68 Hz. See Supplementary Materials Figure 9A Data File. (b). Comparison of classical Doppler effect on *f* changes sensed by equidistant electrodes (same as in (a) above) of a moving source ***S***_***c***_ (same ***v***_***s***_ = 1 cm/s) that emits waves propagating centrifugally away at *π* cm/s (same as spinning spiral source above if not moving in x-direction) utilizing Eq. 21. For this to occur, with same ***λ*** at *π* cm and same frequency at 5 Hz, the wave propagation velocity is 5*π* cm/s. If the electrode was directly on the ***v***_***s***_ vector path, the approaching *f* = 5.34 Hz, and the receding *f* = 4.70 Hz). Note that from “far” away, greater electrode distance, the WF strike *f* is almost identical to that of a spiral wave at a distance. See Supplementary Materials Figure 9B Data File.

**Figure 10.**
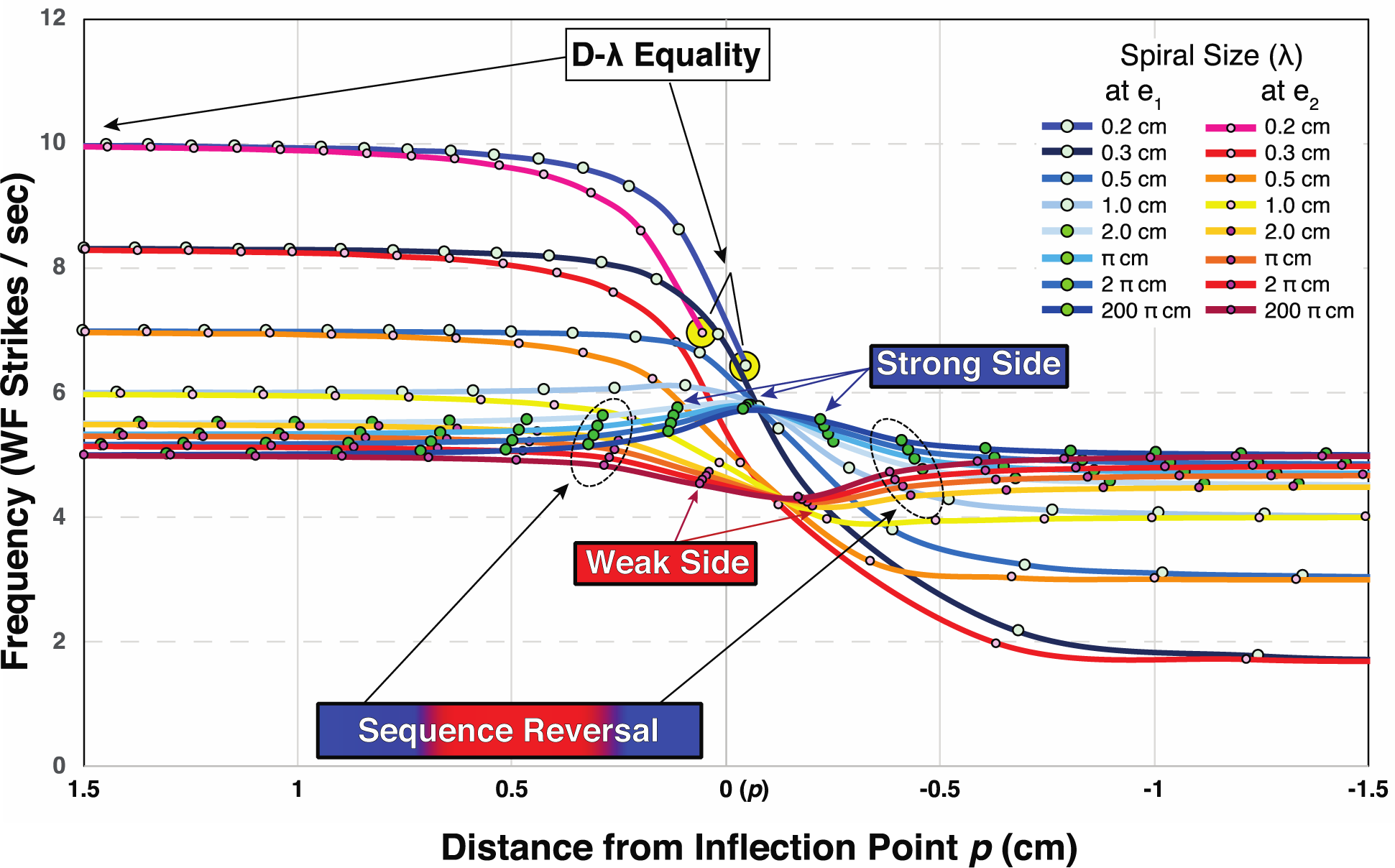
Spiral size on diametric property of SDE. Plots of different spiral sizes vs frequency are shown when electrodes e_1_ and e_2_ (each 2 mm from **p**) are equidistant to ***S***_***Sp***_ along its vector path of motion. ***S***_***Sp***_ moves along the vector with constant ***v***_***s***_ = 1 cm/sec, while spinning at constant ***ω***_***s***_ = 5 Hz, with RDU = 2*π*/0.2cm. Curves are smoothed lines linking the subsequent WF strikes of ***S***_***Sp***_ of same size ***λ***. Pairs of curves show frequency differences between strong side e_1_ (light or dark green dots) and weak side e_2_ (light or dark pink dots). At far distances (x>>r), either approaching or receding, the WF strikes at e_1_ and e_2_ from the largest spiral size (200 *π* cm) have values near ***ω***_***s***_. The WF crest at large ***λ*** approaches that of a radial arm and results approach those similar to a ***S***_***R***_ (see Fig 5 with ***v***_***s***_ = 1 cm/sec, with constant ***ω***_***s***_ = 5 Hz). As ***λ*** gets smaller, the approaching *f* is faster, and receding *f* is slower. Strong side electrode has faster *f* than weak side electrode along the vector path, with the greatest difference observed at the inflection point ***p***. A skewed-diamond pattern emerges at the inflection point with the *f* separation at ***p***. When ***λ*** = D at 0.2 cm, the approaching WF *f* is 10Hz, while at the inflection point WF *f* goes to zero (last WF strikes at yellow circles). An unpaired WF strike on the strong side (3 dark blue arrows) with that on the weak side (2 red arrows) results in the final step before sequence reversal is complete (dashed ovals). See Supplementary Materials Figure 10 Data.

### B. Spiral wavelength effect on diametric property of SDE, λ/D ratio, and surfing

The diametric property of the SDE was shown to be amplified by the closest equidistant electrodes to the vector path ***v***_**s**_ of ***S***_***Sp***_. Utilizing that observation, further assessment of the diametric property was made by varying the spiral size ***λ*** as the ***S***_***Sp***_ moved between electrodes spaced only 2 mm on either side of ***v***_**s**_ (Fig 10). All spiral sizes exhibited the diametric property with strong and weak sides showing their greatest difference in frequencies at the inflection point (skewed diamond-shaped separation at center, center at ***ω***_***s***_). The peak of WF frequencies occur closer to ***p*** than the nadir of weak-side frequencies, that result in the skewed-diamond pattern. The later weak side nadir is a result of the unpaired strong side WF and the sequence reversal. The largest spiral sizes approached frequency changes most similar to RDE seen with rotational wave, where approaching and receding frequencies were equal at relatively long distances from ***p*** with a frequency of ***ω***_***s***_. This result is intuitive since the WF crest of a ***S***_***Sp***_ at the larger ***λ*** (***λ***/D > 1), the frequency would progressively appear more like a radial arm extending out from the center and is expected to resemble the frequency changes of a perfect rotating radial arm as seen in Fig 5. At the other extreme, decreasing ***λ***, resulted in a larger spiral-size component in the slope of the summation angle, *ψ*_***SA***_ of Eq. 31 and Eq. 38, causing greater variation between the approaching and receding ***S***_***Sp***_. In the special situation where ***λ*** = D, the approaching frequency is twice that of ***ω***_***s***_, and the receding frequency is zero. At this surfing ratio, there are no WF strikes because the WF crests continuously straddles the electrodes. At smaller ratio, where ***λ***/D < 1, then WF sequence does not reverse when ***S***_***Sp***_ recedes from ***p*** (not shown). A ***S***_***Sp***_ at a distance from ***p***, the diametric property of the SDE shows that spiral size ***λ***, the *f* changes as observed by electrodes near the vector path ***v***_***s***_ are close to those of a pure rotational wave, while at small spiral size ***λ***, the *f* changes are nearly that of centrifugal waves in the classical Doppler example. A ***S***_***Sp***_ near the inflection point (within 1-2 ***λ***, strong and weak sides create marked differences in *f*, in some cases further increasing the *f* before a sudden decrease. At each ***λ***, near the inflection point ***p***, an unpaired WF strike occurs on the strong side. The unpaired WF strike at e1 was repeatedly observed experimentally during mapping for rotor activity in human atrial fibrillation. We labelled this phenomenon as the “1/2 cycle drop-off” on the weak side (*2*). Most remarkable, the diametric property also explains the profound result that an approaching ***S***_***Sp***_ can cause an increase in *f* simultaneous to a decrease in *f* when observed by 2 electrodes that are equidistant along the entire ***v***_***s***_ of ***S***_***Sp***_.

## IV. DISCUSSION

Wave fronts emitted from rotational and spiral sources do not follow classic Doppler effect phenomena. In a previous investigation (*2*) of direct human heart endocardial surface recordings during atrial fibrillation, a number of new findings were observed from multiple simultaneous recordings of rotors that breached a perimeter of electrodes. A clear diametric frequency change was observed consistently as a rotor approached and then passed between 2 closely positioned electrodes. One electrode recorded an increasing frequency while the second electrode simultaneously recorded a decreasing frequency. These diametric phenomena must occur as the sequence of activation between the 2 electrodes reverses when the rotor passes between them. In addition, an unpaired WF strike occurred at the electrode recording the higher frequency that we had labelled as the ½ cycle drop-off. This paper derives new Doppler equations needed to predict WF strikes frequency of moving and spinning sources of WFs. These equations provide new insight into description of atrial fibrillation, methods of mapping, ablation target planning, in addition to a better understanding of natural phenomena that occurs as a result of rotational and spiral WFs.

Using a trigonometric approach, a new rotational Doppler effect, RDE equation was derived from the creation a new unit, RDU, the family of RDU lines, and their intersection of polar plots of arccotangent curves of the LOS angle ***θ*** between the electrode and the vector path ***v***_***s***_. WF strikes from ***S***_***R***_ occurred only when the ***ψ***_***c***_ equaled the LOS ***θ***. The RDE required a method to identify weak and strong sides of the vector path ***v***_***s***_. A 3-point sequence rule or WF vector method has been provided. The derivation of the RDE provided the capability to then derive the spiral Doppler effect, SDE equation. The spiral WF required a new spiral math to be developed that could plot a spiral summation angle, ***ψ***_***SA***_ as a function of the spiral center’s distance to the inflection point on its path while passing by the observer. WF strikes from ***S***_***Sp***_ occur only when the ***ψ***_***SA***_ equaled the LOS angle ***θ***. Both ***S***_***R***_ and ***S***_***Sp***_ exhibit side-dependent diametric frequency changes (Fig 5 and Fig 8). Both showed a strong side unpaired WF strike at the inflection point as a sequence reversal occurred. Both show staggered potential shear and stall zones on strong and weak sides respectively, each separated by the distance D of the RDU. In both RDE and SDE, observation positions closer to the vector path ***v***_***s***_ of either ***S***_***R***_ or ***S***_***Sp***_ exhibited more profound frequency differences between strong and weak sides.

The SDE shows a transition from WF frequency changes similar to RDE when observing closest to the center of rotation, within 1 ***λ***, to a CDE of frequency changes when observed at distances > 1***λ***. In human recordings during atrial fibrillation (*2*), we had the advantage to record very close (electrode spacing 2mm apart) to cardiac spiral WF sources that have relatively large ***λ***. Figure 9A and B shows the importance of the observational distance. Our ability to detect the diametric property from spiral waves shows that the cardiac rotor sizes that we recorded from were within ranges previously reported (*7, 28, 30*). The closer an observer is positioned to a ***S***_***Sp***_, the greater the differences in timing and thus frequency change observed.

The RDE and SDE equations successfully predict experimental observations of the diametric frequency changes, sequence reversal, and an unpaired strong side WF strike as that found with meandering rotors during atrial fibrillation. In addition, the equations provide further ability to predict phenomena that might be proved experimentally or has been previously observed in other rotational or spiral WF phenomena throughout nature. In recognition to Doppler, the name to describe frequency changes as a result of a moving source is kept, but one must subcategorize them depending upon wave shape. The consistent phenomena observed with rotational and spiral sources that are distinct from classical Doppler effect can be summarized into a descriptive diametric property of RDE and SDE.

### A. The diametric property of RDE and SDE – Rotational Relativity

A general understanding of the diametric property is that different frequencies are recorded that are side-dependent from a moving body that emits a WF that spins about its center at a constant ***ω***_***s***_. An actively spinning WF source at rest in 2D or 3D space creates no differences in frequency recorded from stationary observers anywhere in that 2D or 3D space. All observers record the same exact frequency as that of source’s ***ω***_***s***_ whether it is a ***S***_***R***_ or a ***S***_***Sp***_. The exact time of when the WF strikes each observer may be different as the WF sweeps around 2D or 3D space, but each records a local constant frequency equal to the ***ω***_***s***_ of the source. The RDE and SDE were derived from 2D space, but can be expanded to 3D space, not discussed here. In the 2D derivation of RDE and SDE in this paper, any relative movement between source and observer creates a strong and weak side. The change from resting positions to relative movement between the source and observer immediately results in the separation of 2D space into a strong and weak side half planes. The line of separation of planes is the relative ***v***_***s***_ of the source. The sequence of activation compared to the forward vector path determines whether an observer sits on the strong or weak side.

Figure 11(a) provides a rather simple illustration in which Person R is 100 meters from 2 stationary observers S and W, positioned close together at 2 feet apart. Person B and C are told that they can turn their heads to keep their eyes focused on Person A. Person A moves at a constant velocity directly between persons S and W to a position 100 meters past them while spinning one half of a complete rotation. Person R turns at a constant ***ω***_***s***_, keeping his or her face towards observer W, such that the nose of R and W almost touch at the midpoint of A’s path. Person R completes ½ of a turn when getting 100 meters passed, now facing both S and W again. Person W saw no turning of W at all, while person S saw essentially one full turn. Person S directly saw Person R’s left side, back, right side and front again. Now, in a second pass, person R makes 1½ turns, spinning again in the same direction, moving that same 200-meter distance on a path between observers S and W. At the inflection point directly between S and W, person B sees R’s back, while person S and R are facing each other. Person R continues to move past, completing another ¾ turn facing both observers S and W. Person R made 1 ½ turns, while W observe 1 full turn, while S observed 2 full turns. No matter the number of turns by R, no matter the velocity, nor angular velocity, observer W sees that R turns one less time than observer S. Conservation of momentum is maintained with a strong side WF higher frequency that is met with an equal but opposite weak side WF slower frequency. The most intriguing aspect of the opposite relative changes in frequency between strong and weak sides are all because of just one greater number of rotations sensed between the 2 sides. The side-dependence of frequency observed is rotational relativity. This one rotation difference occurs as the source changes at the inflection point from an approaching S to that of a receding S, when reversal of sequence between strong and weak side occurs. Rotational relativity is independent of n, ***ω***_***s***_, and ***v***_***s***_, source or observer movement, and size. It cannot be understated that a general conclusion of the diametric property of RDE and SDE is that side and sequence matters.

**Figure 11.**
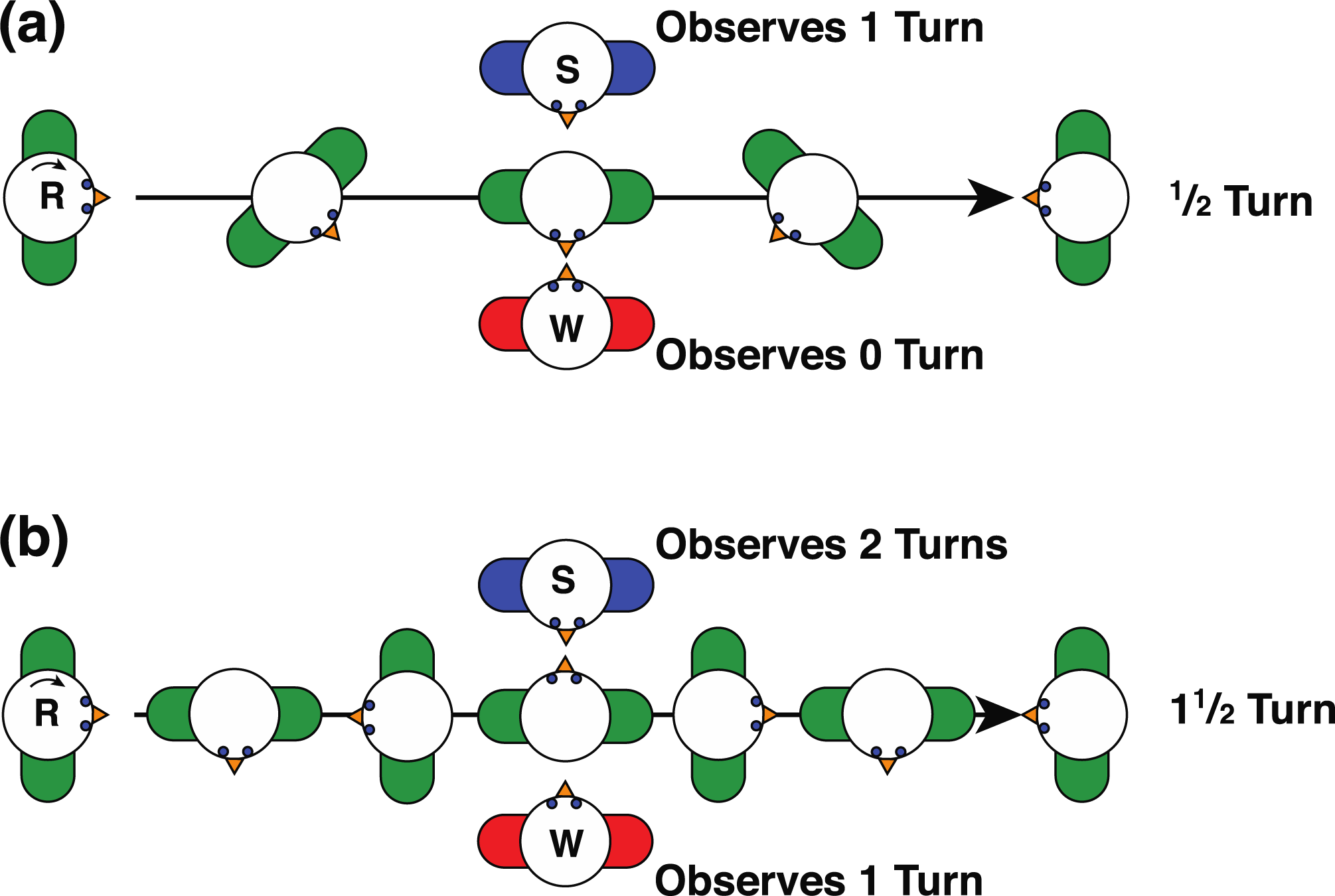
Rotational Relativity. (a) Person R moves along a path between persons S and W. Initially Person R faces both S and W. Person R spins CW making ¼ turn as R reaches midpoint between S and W, facing W. Person R continues on path making another ¼ turn facing both S and W. Person W observed 0 turns since person R faced person W along its entire path. Person S observed 1 turn, observing R’s right side, then back side at the inflection point between S and W, then R’s left side, and back to observing R’s front. (b) Person R moves along a path between persons S and W with a faster spin. Person R spins making ¾ turn as R reaches midpoint between S and W, facing W. Person R continues on path making another ¾ turn. Person S observed 2 turns. Person W observed 1 turn.

If a spiral wave’s energy pushes mass, then it is the diametric property of the RDE and SDE may likely be responsible for spiral growth of crystal formation at the molecular level (*32*), while it provides for the underlying physics a hurricane’s strong and weak sides at a macro weather phenomena level (*33*). In these examples, one can consider the spiral shape is made up of numerous spiral arms sharing the same center. Below the water surface, the strong side of the hurricane has a 3-dimensional build-up of water mass, called an Ekman transport, while above the surface the hurricane has its highest wind speeds. The typical CCW hurricanes hitting the U.S. move North from the Caribbean has its highest wind speeds and storm surge on the eastern side of it eye wall. Investigation of the shear zone identified with the diametric property (as seen in Fig 4) may be worthwhile for future investigations for the possible causes of strong side spawning of 2^nd^ order cyclones (*34, 35*). The 2^nd^ order cyclones, or vortex shedding is evident in studies of Reynold’s number, air turbulence and lift in the description of insect and avian flight (*36, 37*). Creation of one’s own spiral wave and surfing between its crests for a means of propulsion appears to be directly related to computations of specific surfing ratio of D and ***λ*** as seen in Fig 10.

### B. Rotational frequency shifts

Rotational frequency shifts have also been identified in studies of light waves. Frequency shifts have been observed from photons emitted, scattered, or absorbed from rotating bodies, prisms (*38-40*) or lenses (*41*), and derived equations from a quantum mechanics or angular momentum perspective (*42,43*). The rotational shift in frequency of a light beam was determined to be maximal if observed at right angle to the plane of rotation (*40*). However, in non-electromagnetic waves, ***S***_***R***_ and ***S***_***Sp***_ have the maximal frequency shift in the same plane of rotation and right angles to the vector path of the source at the inflection point. The maximal frequency for a ***S***_***Sp***_ was not directly in front of the moving source as seen in classical Doppler.

A natural extension of inquiry leads one to ask how the diametric property of rotation affects the original Doppler conclusions? This can be intuitively answered by creating an imaginary example that two Christian Dopplers existed, each on 2 different planets that were equidistant to a rotating binary star system. Here, a smaller star rotates at a constant ***ω***_***s***_ around a larger star in the same plane of that created by the star system and 2 planets. The binary star system moves on a vector path between the two Christian Dopplers (Doppler 1 and Doppler 2). The vector path separates the plane into two halves. Doppler 1’s planet sits on strong side, while Doppler 2’s planet sits on weak side of the rotation. The maximal highest frequency recorded from each Doppler would occur each time that the smaller star’s tangential path was equal to the LOS to that particular Doppler. The tangential path is always perpendicular to a radial arm from the center of larger star. That radial arm is equivalent to the clock hand of Fig 4. Therefore, instead of recording the time of when the clock hand angle is equal to the LOS, the peak frequency would occur at time ¼ of a rotation (*π*/2 radians) earlier. Simply stated, the peak frequency recorded with each rotation around the larger star would result in the RDU lines shifted by *π*/2 for each Doppler observer. The two Christian Dopplers would have witnessed classic Doppler effects with same peak and nadir frequency achieved. Yet the exact times at which the peak frequency occurred at Doppler 1 and Doppler 2 would be different as well as the subsequent pattern of frequency peaks occurring as the binary star continued on its path. Strong side Doppler 1 would record the peak frequency progressively earlier in time as the binary source approached, while weak side Doppler 2 recorded it progressively later. Once the binary star system passes the inflection point, Doppler 2 would have witnessed one less cycle of frequency shifts. However, the 2 Dopplers would have recorded diametric trends of when the peaks occurred and how many were recorded. Had they looked at trends of the peaks and how many, the 2 Dopplers having opposite results might have developed opposite conclusions and their theories would not match if they didn’t know their twin existed on the opposite side of the binary star. Once again, sequence matters.

### C. Atrial fibrillation: Wave and Source Duality - The Slide(s)-Pivot(s)-Slide(s)/Triggered Model

The diametric property of SDE allows one to slightly modify description of atrial fibrillation that provides new and or modified avenues of treatment strategies to improve outcomes. Action potential WFs in the human heart can only propagate in one direction short distances given the heart’s limited size. All AP WFs, whether propagating outward centrifugally, or as a spiral arm, must encounter lateral boundaries on either side. The WF continues to propagate forward, unless of course the boundaries meet in front, extinguishing the WF. In atrial fibrillation, these physical and/or functional transient lateral boundaries on either side can be considered slits of continuous varying widths. The WF interacts with other WFs, creating lateral boundaries for them as well. The lateral curvature of WF as it emerges past a boundary edge can become a site of vortex shedding creating 2^nd^ order spiral wave rotor source (*28, 45, 45*), same as 2^nd^ order cyclones as discussed above. Jalife et al. put forth a testable dominant frequency hypothesis requiring rotors as the primary drivers of AF with subsequent daughter wavelets and 2^nd^ order rotors through vortex shedding (*46*) to sustain the arrhythmia.

The majority of rotors, spiral waves that were found in our study (*2*) as well as by others (*46,47*), were not sustained more than a few seconds. Our methods had selected 5 rotations or more (typically lasting at least 1 second). Even if a WF pivoted with just 1 or 2 rotations (consider naming this as a swirl), great timing of tissue activation or instantaneous frequency shift of adjacent tissue would be manifest due to the diametric property. One can consider these transient barriers as a sliding pivot for the WF. The length of the slide is dependent upon the length of the barrier. The changing borders of refractory tissue provide new barrier end points from which the WF can bend around, pivot, propagate, slide and may even shed new ***S***_***Sp***_. Focal ectopic source of APs, either from spontaneous or from triggered automaticity, would add to additional drivers or functional barriers of conduction to sustain the duration of AF (*48*). It matters not from what source the AP WF propagates from, as it passes permanent or transient refractory barriers, or slits, it will expand out with curvature at it sides. New pivot points, swirls, or rotors can continue. The emergence of new swirl or spiral source, exhibits a wave/source duality. Just the ability to pivot to unexcited heart tissue faster than propagation of WFs from the sinus node pacemaker cells may be more important than high dominant frequency of transient rotors or swirls. Ablation that targets a specific rotor sites based upon only the highest dominant frequency might fail, since most portions of atrium may change function as a slide, a pivot, swirl, rotor, or a triggered site at different times and conditions. In addition, if one uses the previous assumption that rotors follow CDE and targets tissue based upon location highest dominant frequency (HDF, 5, *31*), one would focus the ablation site to a position directly behind the site of HDF. Yet, the diametric property of SDE would suggest the HDF would not be directly along the rotor’s path, it would be recorded along the strong side. The actual path of a rotor would be a line separating highest frequency and lowest frequency. The strategic goal for treatment of AF remains to straighten WF path, maximize the slide, minimize that pivots, swirls, rotors, or triggered activity with minimal tissue loss. It was recognized early on in the study of rotating sources (*8, 49*), that the drifting of its core created irregularity in the electrical activation. A stationary rotational source of activation would cause a constant frequency of activation, we now know that the motion of the source creates the irregularity observed by the newly identified diametric property of a spiral Doppler effect. A better understanding of frequency changes as a result from moving and spinning WF sources should provide a more comprehensive approach to treatment of AF that might require modification of ablation targets and/or medication customized to a specific patient.

### D. Spiral Doppler effect as phenomena at all levels of nature

Future investigations of the diametric property, RDE, and SDE, might include improving chemical reactions, enantiomer formation, more accurate descriptions and understanding of air turbulence, flight, water eddies, Reynold’s number, weather models, and plotting paths of space travel. The diametric property is a new principle of rotational and spiral wave fronts. The diametric property of rotational and spiral Doppler effects appears to be the critical link between many phenomena at all sizes of nature.

## Supporting information

Figure 4 Data and Web link

Figure 5 Data and Web link

Figure 8 Data and Web link

Figure 9A Data and Web link

Figure 10 Data and Web link

## Acknowledgements

I gratefully acknowledge the support, multiple discussions and identification of additional references by Mason Rubenstein.

## Supplemental Materials

Figure 4 Data and weblink

Figure 5 Data and weblink

Figure 8 Data and weblink

Figure 9A Data and weblink

Figure 9B Data

Figure 10 Data and weblink

